# Relaxin Modulates the Genomic Actions and Biological Effects of Estrogen in the Myometrium

**DOI:** 10.1101/2024.04.15.589654

**Authors:** Sudeshna Tripathy, Anusha Nagari, Shu-Ping Chiu, Tulip Nandu, Cristel V. Camacho, Mala Mahendroo, W. Lee Kraus

## Abstract

Estradiol (E2) and relaxin (Rln) are steroid and polypeptide hormones, respectively, with important roles in the female reproductive tract, including myometrium. Some actions of Rln, which are mediated by its membrane receptor RXFP1, require or are augmented by E2 signaling through its cognate nuclear steroid receptor, estrogen receptor alpha (ERα). In contrast, other actions of Rln act in opposition to the effects of E2. Here we explored the molecular and genomic mechanisms that underlie the functional interplay between E2 and Rln in the myometrium. We used both ovariectomized female mice and immortalized human myometrial cells expressing wild-type or mutant ERα (hTERT-HM-ERα cells). Our results indicate that Rln modulates the genomic actions and biological effects of estrogen in the myometrium and myometrial cells by reducing phosphorylation of ERα on serine 118 (S118), as well as by reducing the E2-dependent binding of ERα across the genome. These effects were associated with changes in the hormone-regulated transcriptome, including a decrease in the E2-dependent expression of some genes and enhanced expression of others. The inhibitory effects of Rln cotreatment on the E2-dependent phosphorylation of ERα required the nuclear dual-specificity phosphatases DUSP1 and DUSP5. Moreover, the inhibitory effects of Rln were reflected in a concomitant inhibition of the E2-dependent contraction of myometrial cells. Collectively, our results identify a pathway that integrates Rln/RXFP1 and E2/ERα signaling, resulting in a convergence of membrane and nuclear signaling pathways to control genomic and biological outcomes.

## Introduction

The uterus must provide and maintain an environment that is conducive to the growth and development of the fetus throughout pregnancy, yet also gradually prepare for parturition, the process of labor and delivery of the fetus at term. Uterine myometrial smooth muscle cell (SMC) plasticity ensures transition between the non-contractile and contractile states in response to physiological signaling cues, at term (1,2). Misregulation of myometrium function can result in numerous obstetrical complications, including preterm birth or Cesarean sections (3,4).

Maintenance of myometrial quiescence and the transition to contractility is regulated through hormones such as progesterone (P4), estradiol (E2), and relaxin (Rln) (5,6). As the actions of P4 decline in late pregnancy, the actions and levels of E2 are enhanced (7,8). The effects of E2 are mediated through nuclear estrogen receptors (ERs) (9,10). ERα is generally expressed at higher levels than ERβ and plays a dominant role in the biology of the female reproductive tract (9,11). The effects of E2 in the myometrium include increased responsiveness to contraction-promoting molecules (e.g., oxytocin and prostaglandins), increased gap junction formation, decrease nitric oxide activity, and increased Ca^+2^ influx (12–14). The peptide hormone Rln has numerous roles in the physiology of pregnancy in humans and other species (15–17). Rln, which is secreted from the corpus luteum and the placenta, mediates its actions through a G-protein coupled receptor, relaxin family peptide receptor 1 (RXFP1) (16). Although Rln is best known for its actions in the cervix (18,19), numerous studies have uncovered diverse roles for Rln in the myometrium (16,17,20,21).

In the uterus, Rln induces uterine growth and development, inhibits apoptosis, promotes smooth muscle cell relaxation via cAMP induction, and promotes vasodilation of the uterine arteries (15,17,22). Pharmacological doses of Rln diminish myometrial contraction induced by either oxytocin or prostaglandins (15,23). A critical role of Rln signaling during parturition is supported by incomplete cervical remodeling and parturition defects in mice with targeted mutation in the *Rln1* (Rln) or *Rxfp1* (Rln receptor) genes (24,25). Interestingly, many of the effects of Rln on reproductive tissues, such as cell proliferation, ECM remodeling, and angiogenic actions, require or are augmented by E2/ERα signaling (16,26,27). Conversely, in some cases, the two hormones have opposing actions. For example, Rln promotes smooth muscle quiescence in the myometrium, while E2/ERα promotes contraction (2,28–30). The molecular mechanisms by which Rln/RXFP1 signaling can promote or oppose the actions of E2/ERα have not been defined.

Many of the biological actions of estrogens are mediated by ERα, which functions primarily as a ligand-regulated, DNA-binding transcription factor (TF) in the nuclei of estrogen-responsive cells. Nuclear ERα dimerizes in response to E2 and binds to many thousands of ERα binding sites (ERBSs) across the genome, collectively called the ERα ‘cistrome’ (31–35). Most ERα binding sites are pre-bound by “pioneer” transcription factors (e.g., FoxA1, AP-2γ) prior to estrogen exposure, which facilitate the binding of ERα to a repertoire of predetermined sites (36,37). Many ERα binding sites contain a DNA sequence motif called the estrogen response element (ERE) (10). The binding of ERα to genomic DNA promotes the coordinated recruitment of coregulator proteins that establish an active ‘enhancer,’ leading to chromatin looping and target gene transcription (38–42).

Importantly, ERα actions can be modulated by post-translational modifications (PTMs). For example, ERα can be phosphorylated on multiple sites by a wide range of kinases, which regulate various functions such as hormone sensitivity, nuclear localization, DNA binding, protein and chromatin interactions, and gene transcription (43,44). Phosphorylation at serine 118 has been reported to be important for ERα dimerization, interactions with coactivators such as CBP/p300 and SRC, receptor activation, and ERα enhancer formation (45–47). Although serine 118 phosphorylation may promote the progression of uterine leiomyomas, its effects on myometrium function have not been characterized (48,49).

In the studies described herein, we examine the extent and nature of functional interactions between the E2 and Rln signaling pathways in the myometrium. In addition, we delineate the mechanisms underlying functional interplay between the E2/ERα and Rln/RXFP1 signaling pathways at the molecular and physiological levels in myometrial tissue and cells during quiescence and contractility.

## Methods and Materials

### Antibodies, hormones, and inhibitors

The antibodies used were as follows: ERα, rabbit polyclonal generated in house (50); phospho-ERα (S118), mouse monoclonal (Cell Signaling Technology, 2511S; RRID: AB_331289); phospho-ERα (S104/106) (Abcam, ab75753; RRID: AB_1310197); phospho-ERα (S167), rabbit monoclonal (Cell Signaling Technology, 64508; RRID: AB_2799660); H3K27ac, rabbit polyclonal (Abcam, ab4729; RRID: AB_2118291); H3K4me1, rabbit polyclonal (Abcam, ab8895; RRID: AB_306847); ERK1/2 (Cell Signaling Technology, 4695; RRID: AB_390779); phospho-ERK1/2 (Cell Signaling Technology, 4370; RRID: AB_2315112); DUSP1, rabbit polyclonal (Cell Signaling Technology, 35217; RRID: AB_3371713); DUSP5, rabbit polyclonal (Thermo Fisher Scientific, PA5-85961; RRID: AB_2802762); β-actin, mouse monoclonal (Cell Signaling Technology, 3700; RRID: AB_2242334); goat anti-rabbit IgG (H+L)-HRP conjugate (BioRad, 1706515; RRID: AB_11125142); and goat anti-mouse IgG (H+L)-HRP conjugate (BioRad, 1706516; RRID: AB_2921252). The hormones used were as follows: 17β-estradiol (E2; Sigma) and recombinant human relaxin-2 (Rln; Peprotech). The MEK inhibitor Trametinib (GSK1120212) (51) was from MedChemExpress (HY-10999).

### Mouse models, ovariectomy, and hormone administration

All animal studies were conducted in accordance with the standards of humane animal care as described in the NIH Guide for the Care and Use of Laboratory Animals. The research protocols were approved by the IACUC office at the University of Texas Southwestern Medical Center. The mouse colony was maintained in a barrier facility and the health status of the mice was assessed on a quarterly basis. We used the following mouse strains: Wild-type (*Rxfp1^+/+^*) and relaxin receptor knockout (*Rxfp1^-/-^*) (52) on a C57BL/6 background housed under a 12 h-light/12 h-dark cycle at 22^◦^C. Female mice at 7 to 8 weeks of age were ovariectomized and rested for 2 weeks to deplete endogenous ovarian hormones. For experiments, the ovariectomized mice were injected i.p. with 0.1 mL of saline (vehicle), 2.5 μg/mL 17β-estradiol (E2, E; Sigma), 80 μg/mL 0.1% BSA-PBS recombinant human relaxin-2 (Rln, R; Peprotech) and a combination of E+R for the specified amount of time (five mice per treatment per biological replicate). Mice were sacrificed under 2,2,2-tribromomethanol (Sigma) anesthesia. The uteri were harvested and further enriched for myometrium by sterile scraping, washing in cold 1x PBS and blotting with a paper towel. The enriched myometrial tissues were flash frozen for subsequent genomic and biochemical assays.

### Cell culture and treatments

Telomerase-immortalized human myometrial (hTERT-HM) cells (53) were cultured in phenol-red free DMEM/F12 (ThermoFisher Scientific) with 10% fetal bovine serum (FBS, v/v), 1% penicillin/streptomycin at 37°C in 5% CO_2_. hTERT-HM cells with doxycycline (Dox)-inducible expression of ERα (wild-type, WT) and ERα mutants (S118A and S118E) were generated by lentivirus-mediated transduction using the pINDUCER20 vector (RRID: Addgene_44012), as described below.

For experiments, the cells were treated with vehicle (DMSO), 100 nM 17β-estradiol (E2, E; Sigma, St. Louis, MO, USA), 0.2 μg/mL recombinant human relaxin-2 in 0.1% BSA-PBS (Rln, R; Peprotech, Rocky Hill, NJ, USA) and a combination of E+R for the specified amount of time. Prior to treatment, the media was supplemented with 5% charcoal-dextran-treated calf serum (CDCS) for 3 days. The cells were treated with doxycycline for 24 hours prior to the different treatments.

### Generation of recombinant ERα cDNAs and expression constructs

Plasmids containing gene of interest, ERα (WT, S118A and S118E), were used as a template for PCR amplification using following primers:

- S118A Forward: 5’-GCAGGAAAGGCGCCAGCTGCGGCGG-3’
- S118A Reverse: 5’-CCGCCGCAGCTGGCGCCTTTCCTGC-3’
- S118E Forward: 5’-CTGCAGGAAAGGCTCCAGCTGCGGCGGC-3’
- S118E Reverse: 5’GCCGCCGCAGCTGGAGCCTTCCTGCAG-3’

PCR-amplified fragments were cloned into pINDUCER20 using a ligation-independent cloning protocol with 5x In-Fusion HD Enzyme Premix (Takara Bio USA, Inc.) per manufacturer’s protocol. The recombinant constructs were confirmed by DNA sequencing.

### Generation of hTERT-HM cells stably expressing ERα

hTERT-HM cells with Dox-inducible expression of ERα (WT, S118A and S118E) were generated by lentivirus-mediated transduction. The lentiviral ERα expression constructs or the empty vector were co-transfected with additional virus constructs (pRSV-Rev, pMDLg, pCMV-VSV-G) into HEK-293 FT cells using Polyjet reagent (SignaGen Laboratories). The medium containing the transfection reagent was replaced with fresh medium (DMEM) 7 hours post-transfection. The lentivirus containing supernatant was collected and filtered for transduction. hTERT-HM cells were infected with recombinant lentivirus at a multiplicity of infection (MOI) or approximately 1. One day after infection, the cells were allowed to recover in fresh medium for 24 hours and were then subjected to drug selection using G418. The expression of ERα under Dox induction was confirmed by Western blotting.

### Knockdown of *DUSP1* and *DUSP5* mRNAs using siRNAs

Commercially available siRNAs targeting *DUSP1* mRNA (Sigma, SASI_Hs01_00098748) and *DUSP5* mRNA (Sigma, SASI_Hs02_00337568) were transfected into hTERT-HM-ERα cells at a final concentration of 30 pmol using Lipofectamine RNAi MAX reagent (Invitrogen, 13778-150) according to the manufacturer’s instructions. All experiments were performed 72 hours after siRNA transfection.

### Western blotting

For determination of protein levels and phosphorylation by Western blotting, extracts were prepared from mouse myometrial tissue or cultured hTERT-HM-ERα cells from a minimum of three biological replicates. Samples were collected for each of the four treatment time points (vehicle, E2, Rln, and E+R) from ovariectomized mice or hTERT-HM cells (1 hour and 30 min treatment times, respectively). Frozen myometrial tissue was powdered using a metal pulverizer. Whole cell lysates of the powdered myometrial tissues and hTERT-HM-ERα grown on 10 cm diameter plates were prepared in ice cold Whole Cell Extract Buffer (10 mM Tris-HCl pH 7.5, 0.5 M NaCl, 2.5 mM EDTA, 1% Nonidet P-40), supplemented with Halt Protease and Phosphatase Inhibitors (Thermo Scientific). The cells were resuspended by pipetting, lysed, and incubated on ice for 30 min. The lysates were centrifuged at maximum speed in a microcentrifuge for 10 min at 4°C and the supernatants were collected as the whole cell extracts. The extracts were subjected to colorimetric bicinchoninic acid (BCA) protein assay (Pierce). Fifteen to 20 μg of total protein were analyzed on 8% polyacrylamide-SDS gel and transferred to a nitrocellulose membrane. Membranes were blocked at room temperature for 1 hour in 5% nonfat dry milk (blocking grade, Bio-Rad) dissolved in Tris-buffered saline containing 0.1% Tween 20 (TBST). The blots were incubated with primary antibody (listed above) in blocking solution overnight at 4°C, washed in TBST, and incubated with horseradish peroxidase (HRP)-labeled secondary antibodies for 1 hour at room temperature. The membrane was imaged with a Bio-Rad ChemiDoc MP Imaging System (BioRad).

### Isolation of total RNA

Total RNA was isolated from myometrial tissue or cultured hTERT-HM-ERα cells using an RNeasy Plus Mini Kit (Qiagen), according to manufacturer’s instructions. Myometrial tissue was homogenized in RLT buffer with a cooled blade homogenizer (Power Gen25; Fisher). hTERT-HM-ERα cells were collected and lysed in RLT buffer. The samples were centrifuged to remove debris and passed through a gDNA eliminator column, followed by RNA binding column. After multiple washes, total RNA was eluted with RNase-free water. The eluted RNA was assayed for quality by electrophoresis (Agilent Technologies, Santa Clara, CA, USA). Only samples with RIN values greater than 8 were included in subsequent analyses.

### Reverse transcription-quantitative PCR (RT-qPCR)

Analysis of the levels of specific mRNAs were assessed by RT-qPCR. Total RNA was reverse transcribed using MMLV reverse transcriptase (Promega, M150B) with oligo(dT) primers (Sigma-Aldrich). The resulting cDNA was analyzed by qPCR using the primer sets listed below using a LightCycler 480 real-time PCR thermocycler (Roche):

- *DUSP1* Forward: 5’-ACCACCACCGTGTTCAACTTC-3’
- *DUSP1* Reverse: 5’-TGGGAGAGGTCGTAATGGGG-3’
- *DUSP5* Forward: 5’-TGTCGTCCTCACCTCGCTA-3’
- *DUSP5* Reverse: 5’-GGGCTCTCTCACTCTCAATCTTC-3’

### RNA-seq library preparation and sequencing

RNA-seq libraries from mouse myometrial tissue and hTERT-HM-ERα cells (WT and S118E) were prepared as follows using a protocol described previously (54). Briefly, biological replicates were generated by pooling 1.5 μg of total RNA from five animals for a total of 7.5 μg of total RNA per replicate. A total of ten animals were used for each treatment time point. Polyadenylated RNA was isolated from total RNA using Oligo(dT)25 Dynabeads (Invitrogen) and reverse transcribed using SuperScript III Reverse Transcriptase (Invitrogen). Second strand synthesis was performed using DNA polymerase I. Complementary DNA ends were repaired, tailed with dATP, and ligated with barcode adapters for Illumina sequencing platform. Ligated libraries were selected for average size 270 bp, PCR amplified for 9 cycles, and sequenced on an Illumina NextSeq 500 in a 75 bp single-end format in the McDermott Center Next Generation Sequencing Core at UT Southwestern Medical Center to a total read count of ∼40 M per condition.

### Analysis of RNA-seq data

#### Quality check, alignment, and differential expression

The raw data were subjected to QC analyses using the FastQC tool (55). The reads were aligned to the mouse genome (mm10) or the human genome (hg38) using TopHat v.2.0.12 (56). Using aligned reads as input, we employed cufflinks v.2.2.1 (57) to assemble the reads into transcripts using Gencode v.M5 annotations. We then used cuffdiff v.2.2.1 (57) to call differentially regulated transcripts. All programs were run with default parameters. The expression of differentially expressed genes was visualized as heatmaps using Java Treeview (58). To categorize treatment-regulated genes, we filtered for genes with: (1) a p-value < 0.01 and fold change cutoff ≥ 2.0 or ≤ 0.5 (log2 ≥ 1.0 or ≤ -1.0) for mouse myometrial cells and (1) a p-value < 0.01 and fold change cutoff ≥ 1.5 or ≤ 0.67 (log2 ≥ 0.58 or ≤ -0.79) for hTERT-HM-ERα cells.

#### Gene ontology (GO) analyses

Genes associated with ERα peaks depleted by Rln treatment were evaluated with WEB-based GEne SeT AnaLysis Toolkit (WebGestalt) (59). Multiple gene ontology terms relating to collagen metabolism and extracellular matrix (ECM) organization were identified. Additional gene and pathways analyses were performed using Gene Ontology (GO) annotations (60) for GO biological process gene sets and the PANTHER enrichment analysis tool (61).

### Chromatin immunoprecipitation (ChIP) in mouse myometrial tissue

#### Tissue preparation and crosslinking

Pools of frozen myometrial tissue (5 myometrium per pool; 60 to 80 mg of tissue) were powdered using a metal pulverizer. The powdered myometrial pool was immediately resuspended in 1% formaldehyde in phosphate-buffered saline (PBS) with a volume of 10 mL per pool, then incubated on a rotating platform at room temperature for 10 min. Crosslinking was quenched by adding 5 mL per pool of 2.5 M glycine, with a subsequent 5 min incubation at room temperature. The crosslinked tissue was collected by centrifugation at 1000 x *g* for 3 min at room temperature. The supernatant was decanted, and the tissue was resuspended in 4°C PBS with added protease inhibitor cocktail (Roche Molecular Biochemicals) at 10 mL per pool, with incubation on ice for 2 min. The samples were centrifuged at 1000 x *g* for 3 min at 4°C. The PBS wash was repeated, and the tissue was resuspended in 2.5 mL per pool of homogenization buffer (50 mM Tris-HCl pH 7.5, 1% Nonidet P-40, 0.25% deoxycholic acid, 1 mM EDTA) supplemented with proteinase inhibitor cocktail.

#### Preparation of soluble chromatin

The crosslinked tissue was homogenized using a cooled blade homogenizer (Power Gen25). The pellet was collected by centrifugation at 1000 x *g* for 5 min at 4°C. The supernatant was decanted and the pellet was resuspended in 1 mL per pool of Farnham Buffer (5 mM PIPES, pH 8.0, 85 mM KCl, 0.5% Nonidet P-40, 1 mM DTT, 0.1 mM PMSF) supplemented with proteinase inhibitor cocktail by mixing five times with a pipettor and then incubated on ice for 10 min. The chromatin was pelleted by centrifugation for 1000 x *g* for 10 min at 4°C and resuspended in SDS Lysis Buffer (50 mM Tris-HCl pH 7.9, 10 mM EDTA, 1% SDS, 1 mM DTT) supplemented with proteinase inhibitor cocktail and histone deacetylases inhibitors (10 mM sodium butyrate, 5 mM nicotinamide). Resuspended chromatin was incubated on ice for 10 min, split into 300 μL aliquots, and sonicated on ice using a Bioruptor UCD-200 (Diagenode) on the high setting for 9 cycles, 30 sec on and 30 sec off with 1 min rest, 4 times, to obtain genomic DNA fragments of approximately 200 to 400 bp, as determined by gel electrophoresis. Sonicated chromatin was centrifuged at maximum speed in a microcentrifuge for 1 min at 4°C to remove insoluble debris, and the soluble chromatin supernatant was flash frozen in liquid N_2_ and stored at -80°C until use.

#### Immunoprecipitation (IP)

Chromatin was diluted 1:9 with Dilution Buffer (20 mM Tris-HCl pH 7.9, 300 mM NaCl, 2 mM EDTA, 0.5% Triton X-100, 1 mM DTT, 0.2 mM PMSF) supplemented with proteinase inhibitor cocktail and histone deacetylases inhibitors. The diluted chromatin was pre-cleared with 20 μL/mL protein A-agarose beads (ThermoFisher Scientific; 50% slurry in 10 mM Tris-HCl pH 8.1; 1 mM EDTA) on a nutator for 1 hour at 4°C. The pre-cleared chromatin was centrifuged in a microcentrifuge at 1000 x *g* for 3 min at 4°C. The pre-cleared supernatant was divided into input (2.5%) and a 1 mL aliquot for immunoprecipitation. Immunoprecipitation was performed overnight at 4°C using the polyclonal antibodies for ERα, H3K27ac, or H3K4me1 described above. For a control, rabbit IgG (BioRad; PRABP01) was added at 1:200.

Immune complexes were collected by the addition of 80 μL of protein A-agarose beads (Cell Signaling Technology; 9863) with an additional 2 hour incubation at 4°C with mixing. The samples were centrifuged in a microcentrifuge at 500 x *g* at 4°C for 1 min to collect the resin. The resin was washed four times for 10 min each in Low Salt Wash Buffer (20 mM Tris-HCl pH 7.9, 2 mM EDTA, 125 mM NaCl, 0.05% SDS, 1% Triton X-100), four times for 10 min each in High Salt Wash Buffer (20 mM Tris-HCl pH 7.9, 2 mM EDTA, 500 mM NaCl, 0.05% SDS, 1% Triton X-100), one time with LiCl Wash Buffer (10 mM Tris-HCl pH 7.9, 1 mM EDTA, 250 mM LiCl, 1% Nonidet P-40, 1% sodium deoxycholate), and one time in TE. All buffers were supplemented with proteinase inhibitor cocktail and histone deacetylases inhibitors. The cross-linked chromatin was then eluted twice using Elution Buffer (1% SDS and 100 mM sodium bicarbonate) at room temperature for 15 min. The eluted chromatin was heated at 65°C overnight in Elution Buffer supplemented with 100 mM NaCl to reverse the formaldehyde crosslinking. RNase was added to 3 μg/mL final concentration was added and incubated at 37°C for 30 min, followed by inactivation with Tris-HCl pH 7.9, 5 mM EDTA, 50 μg/mL proteinase K at 55°C for 2 hours. The immunoprecipitated DNA fragments were purified by phenol:chloroform extraction, followed by ethanol precipitation. The purified ChIPed DNA was stored at -20°C until it was analyzed by quantitative real time PCR or ChIP-seq as described below.

### Chromatin immunoprecipitation reverse transcription-quantitative PCR (ChIP-qPCR)

Purified ChIPed DNA from mouse myometrial tissue was analyzed by reverse transcription-quantitative PCR using the following primers:

- *Igf1* Forward: 5’-GACACACACACAGACCCTAAA-3’
- *Igf1* Reverse: 5’-GTTGTGCTGGAGACCAAGT-3’
- *Pgr* Forward: 5’-TTGTAAAGAGGTGGGATAATGCT-3’
- *Pgr* Reverse: 5’-GTCCAAGGGTGTGTGGTAAA-3’
- *Nrip1* Forward: 5’-GTACTCCTCACTGACACCAAATAG-3’
- *Nrip1* Reverse: 5’-CCTTGCTGAGTGTCGTTTCT-3’

### ChIP-seq library preparation and sequencing

ChIP-seq libraries were generated from two biological replicates for each condition as described previously (62,63). Immunoprecipitated (ChIPed) DNA was purified further by polyethylene glycol (PEG)-mediated precipitation (64) using PEG Precipitation Solution (20% in 20 mM Tris-HCl pH 7.9, 4 mM EDTA, 2.5 M NaCl). After purification, 5 ng (for ERα) and 10 ng (for H3K27ac and H3K4me1) of ChIPed DNA for each treatment condition was used to generate libraries for sequencing, as previously described with some minor modifications (63). Briefly, the ChIPed DNA was end-repaired and a single “A”-base overhang was added using the Klenow fragment of *E.coli* DNA polymerase. The A-modified DNA was ligated with Illumina sequencing adapters using the Illumina TruSeq DNA Sample Prep Kit. The ligated DNA (250 ± 25 bp) was size-selected using a bead-based method with magnetic beads (AmPure XP beads, Beckman Coulter) isolate DNA fragment sizes of interest. The size-selected DNA was amplified by PCR (9 to 11 cycles as determined empirically) and purified using AmPure XP beads. The final libraries were run on agarose gel, and extracted using QIAquick gel extraction kit (Qiagen). The final libraries were subjected to QC (size, purity, adapter contamination). The quality of libraries was assessed using a High Sensitivity D1000 screen tape on a 2200 TapeStation (Agilent) and quantified using a Qubit dsDNA HS assay kit (ThermoFisher Scientific). The libraries with unique adapter barcodes were multiplexed and sequenced on an Illumina HiSeq 2500 (single-end, 75 base reads) to a depth of ∼40 M reads per condition.

### Analysis of ChIP-seq data

#### Quality check, alignment, and peak calling

The raw data were subjected to QC analyses using the FastQC tool (55). The ChIP-seq reads were trimmed using Cutadapt v.1.9.1 (65) and mapped to the mouse reference genome (mm10) using Bowtie v.1.0.0 (66). Uniquely mappable reads were converted into bigWig files using BEDTools v.2.25 (67) for visualization in the UCSC genome browser (68). The aligned reads were used to measure library complexity using BEDTools (v 2.17.0) (67) and met minimum ENCODE data quality standards (69). ERα peaks under all the treatment conditions were called versus input as control using HOMER software (70) and an FDR cutoff 0.05.

#### Peak Annotation and clustering

In order to study the effects of the treatments on the ERα cistromes identified by HOMER, the peaks identified upon a single treatment were combined with the peaks identified upon co-treatment (E+R). The peaks that were gained, maintained, and depleted upon co-treatment were identified as described previously (62,71). A cut-off of 2MAD (median absolute deviation) was used for the ERα peaks to categorize them into gained, maintained, and depleted peaks. The normalized read counts in the 2 kb window (± 1 kb) around the peak categories was visualized as boxplots using the box plot function in R. Wilcoxon rank sum tests were performed to determine the statistical significance of all comparisons.

#### Heatmaps

The read densities surrounding the gained, maintained, and lost ERα peaks [5 kb (± 2.5 kb) for ERα and 10 kb (± 5 kb) for H3K27ac and H3K4me1] were determined using the annotatePeaks.pl module of HOMER software (70) and visualized as heatmaps using Java Treeview (58).

### Collagen-based cell contraction assays

Collagen-based cell contraction assays were performed as described previously (53). Briefly, hTERT-HM-ERα cells (WT, S118A and S118E), with or without siRNA-mediated knockdown of *DUSP1* or *DUSP5*, were plated in collagen gel. Subconfluent cells were collected with trypsin, centrifuged at 1,000 x *g* for 5 min, and resuspended in serum-free DMEM/F12 medium. The cells were treated with Dox for 24 hours prior to setting up the assay to induce the expression of ERα. Cells without Dox treatment served as control. The number of cells was quantified, and their viability was ascertained by staining with trypan blue. A solution of type I rat tail collagen (Cultrex rat collagen I, 3440-100-01, R and D Systems) was adjusted to pH 7.2 with 0.1% glacial acetic acid and 1 M NaOH and a final concentration of 3 mg/mL. An amount of hTERT-HM-ERα cells (WT, S118A and S118E) was added to the neutralized collagen solution to achieve 1 x 10^5^ cells per well in a 12-well plate, mixed, and incubated on ice for 5 min. One half mL of collagen gel-cell suspension was plated per well in a 12-well culture plate, with subsequent incubation for 1 hour at 37°C to allow gelling. After incubation, 1 mL of fresh DMEM/F12 medium containing DMSO (vehicle), 100 nM E2, 0.2 μg/mL Rln, or a combination of E+R was added to the wells. After 20 hours, the gel matrices were gently detached from the sides and lifted off the bottom of the well. The area of each gel matrix was measured periodically for up to 48 hr. Images of the floating gels were captured using Bio-Rad ChemiDoc MP Imaging System (Bio-Rad). The area of each gel was determined using in ImageJ (v.1.52k) (72). The percentage of gel contraction per well was calculated as:

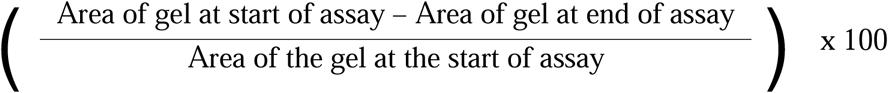

### Data availability

The new RNA-seq and ChIP-seq data sets generated for this study are available from the NCBI’s Gene Expression Omnibus (GEO) database (http://www.ncbi.nlm.nih.gov/geo/) using accession number GSE244843.

## Results

### Rln signaling inhibits the E2-induced genomic binding of ERα in mouse myometrium

To better understand the genomic and molecular mechanisms underlying interplay between the E2/ERα and Rln/RXFP1 signaling pathways in the myometrium, we used 7-to 8-week old ovariectomized female C57BL/6 mice injected i.p. with 0.1 mL of vehicle (V; saline), E2 (E; 2.5 μg/mL), recombinant human relaxin-2 (R, 80 μg/mL) or both (E+R) for 1 or 2 hours [for chromatin immunoprecipitation-sequencing (ChIP-seq) or RNA-sequencing (RNA-seq), respectively] (Fig. 1A). We then collected the myometrium and subjected the tissue to Western blotting for ERα and various genomic analyses. As shown in Fig. 1B, the treatments had no discernable effect on the levels of nuclear ERα by Western blotting.

**Figure 1.**
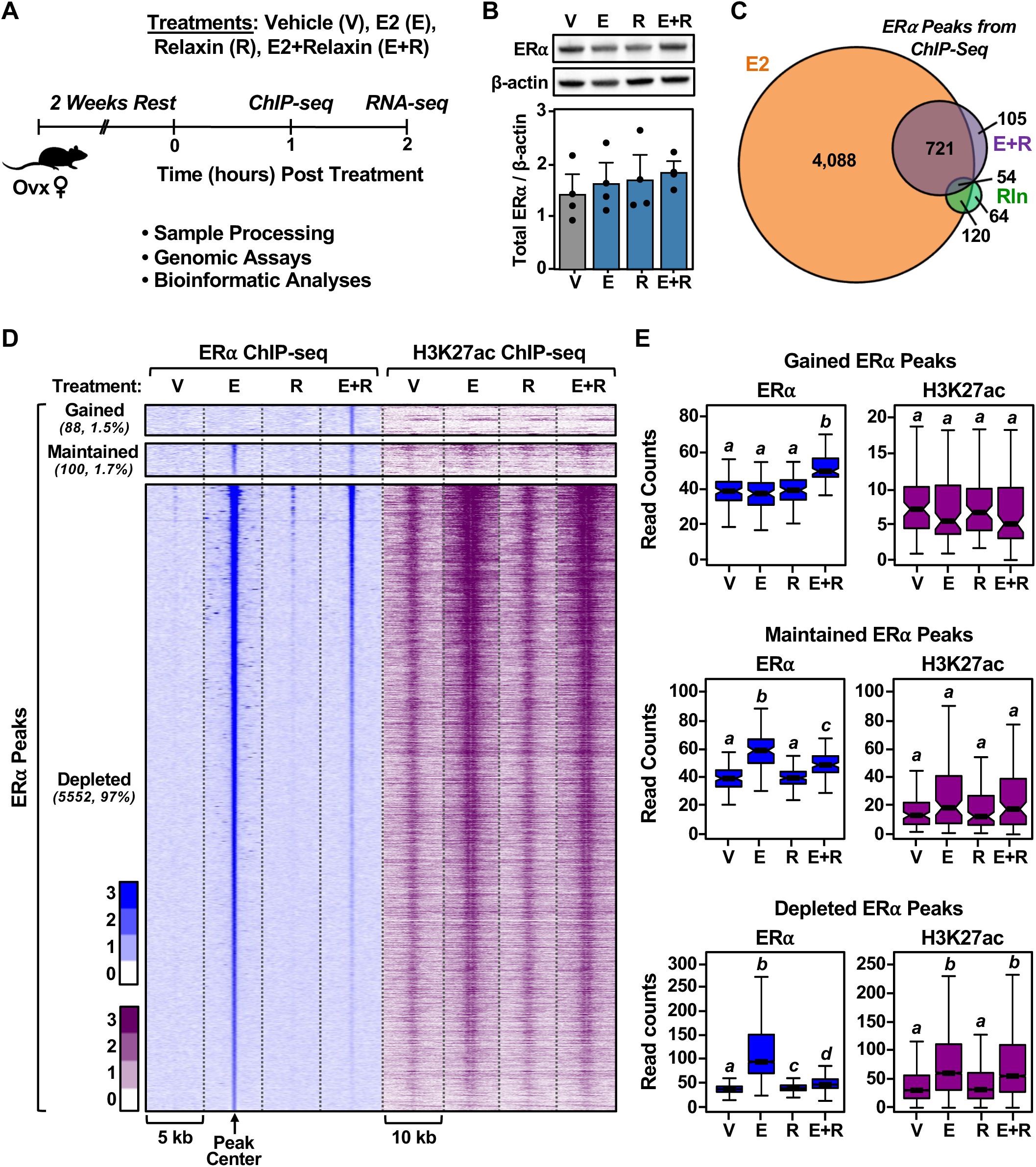
Rln reorganizes the ERα cistrome in the mouse myometrium by promoting the depletion of ERα from its enhancers. **(A)** Experimental schematic for surgical preparation and treatment of female mice, sample processing, and genomic and downstream bioinformatic analyses. C57BL/6 ovariectomized female mice were treated with vehicle (V, 0.1 mL saline), 17β-estradiol (E2; E, 2.5 μg/mL), recombinant human relaxin-2 (R, 80 μg/mL 0.1% BSA-PBS) or both (E+R) for 1 or 2 hours. **(B)** Western blots *(top)* and densitometric quantification *(bottom)* of ERα protein in mouse myometrium samples collected at 1 hour after the indicated treatments. β-actin is a loading control. Each bar represents the mean ± SEM, n = 3. No significant differences were observed. **(C)** Venn diagram showing the number of ERα enhancer peaks under each treatment condition as revealed by ChIP-seq. **(D)** Heatmaps of ERα ChIP-seq data *(left)* from myometrium obtained from mice treated with vehicle (V), E2 (E), Rln (R), or both (E+R). The changes shown in the heatmap were centered relative to the control (vehicle, V) condition. Heatmaps of H3K27ac ChIP-seq data *(right)* at the genomic loci where ERα was bound under the same treatment conditions. **(E)** Box plots of normalized read counts for ERα and H3K27ac ChIP-seq data at three distinct sets of ERα binding sites (gained, maintained, depleted). Bars marked with different letters are significantly different (Wilcoxon rank sum test, p < 0.001).

To explore the effects of Rln signaling on the genomic actions of E2 in the mouse myometrium, we determined the ERα cistrome by ChIP-seq in myometrium isolated from mice subjected to the four treatments listed above. With E2 treatment, we observed a robust induction of ERα binding across the genome as indicated by a large increase in the number of statistically significant ERα binding sites, as expected (4,983 sites; Fig. 1C), as determined by peak calls in ChIP versus input from HOMER (70). Remarkably, we observed a dramatic reduction of E2-induced ERα binding across the genome with Rln cotreatment as indicated by a dramatic reduction in the number of statistically significant E2-induced ERα binding sites (880 sites; Fig. 1C), with 775 sites overlapping those observed in the E2 only condition.

To quantify in more detail the effects of Rln treatment on E2-induced ERα binding, we used read counts and the universe of ERα binding sites to determine median absolute deviation (MAD) using methods described by Rao et al. (71), which allowed us to determine “gained,” “maintained,” and “depleted” ERα binding sites. Again, we observed a dramatic reduction in E2-induced ERα binding sites upon cotreatment with Rln, as shown in a heatmap (Fig. 1D), box plots (Fig. 1E), and metaplots (Supplemental Fig. S1). We also examined the enrichment of two histone modifications associated with enhancer activity, histone H3 lysine 4 monomethylation (H3K4me1) and histone H3 lysine 27 acetylation (H3K27ac) (73,74), at the ERα binding sites. We observed that in spite of the significant reduction in ERα binding at the “depleted” sites upon cotreatment with Rln, significant enrichment of H3K27ac and H3K4me1 was maintained (Figs. 1, D and E; Supplemental Fig. S2). Together, these analyses indicate that Rln signaling broadly inhibits the E2-induced binding of ERα in mouse myometrium from ovariectomized animals. But, the maintenance of H3K4me1 and H3K27ac enrichment indicates that these enhancers may remain open, accessible, and active.

### Rln signaling alters the E2-regulated transcriptome in mouse myometrium

To relate the changes in the E2-dependent ERα cistrome in response to Rln signaling to gene expression outcomes in the mouse myometrium, we first performed RNA-seq under the conditions described above. We expressed the results in a heatmap, Venn diagram, and tabular representation to illustrate differences in gene expression (Fig. 2, A-C). These results demonstrate that E2 is a potent regulator of gene expression in the myometrium, with >1,000 genes upregulated and >1,000 genes downregulated, as expected (Fig. 2C). Interestingly, Rln also regulated the expression of many genes (>400 upregulated and >500 downregulated) (Fig. 2C). Alterations in the E2-regulated transcriptome by cotreatment with Rln were also evident; 372 E2-regulated genes lost significant regulation and 673 E2-regulated genes gained significant regulation upon cotreatment with Rln (Fig. 2, A and B). The response of these gene sets in aggregate to the various treatments are evident in quantitative box plots (Fig. 2D).

**Figure 2.**
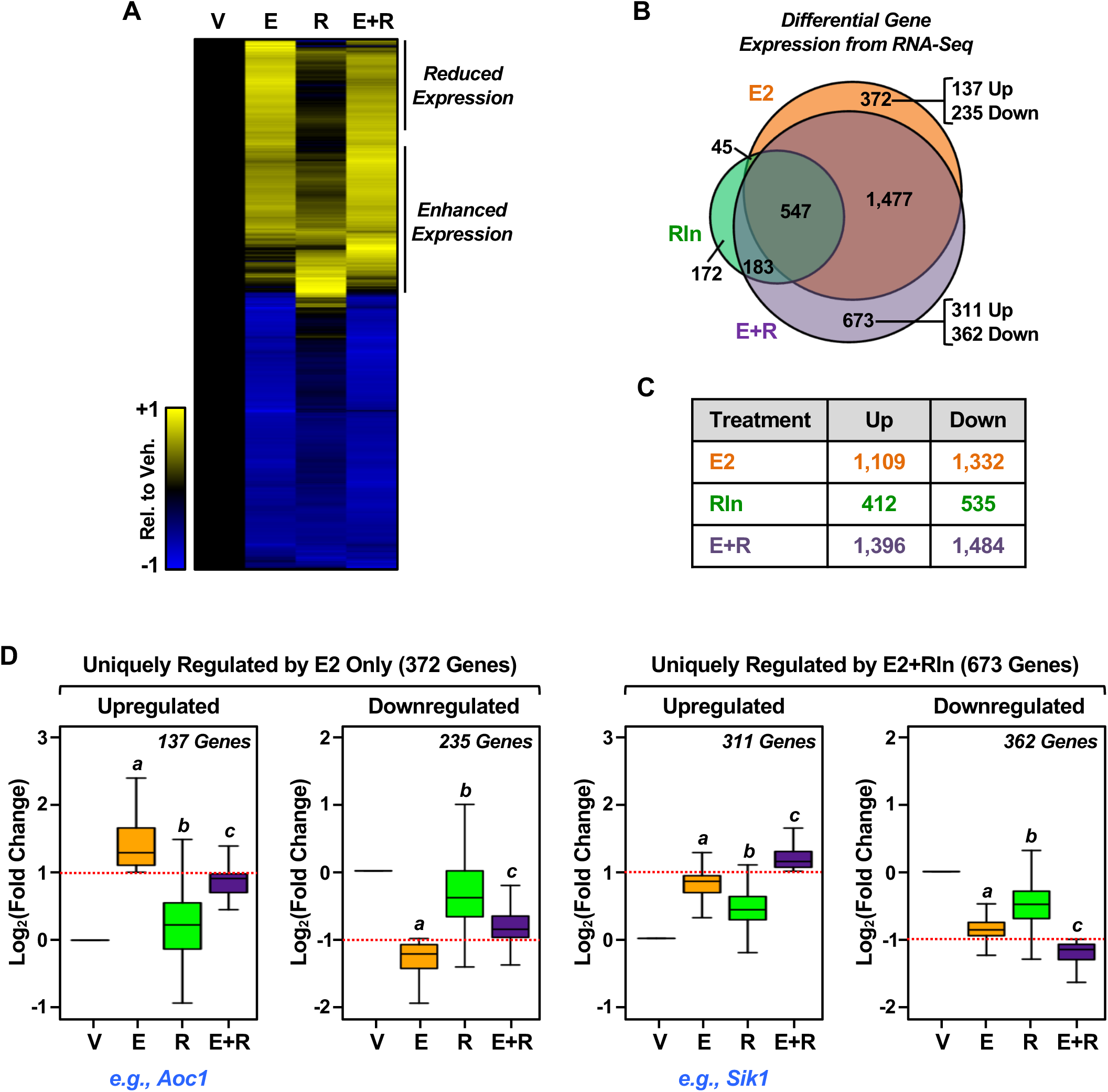
Genomic analyses of the transcriptional response to E2 and Rln mouse myometrium. **(A through C)** Three representations of the RNA-seq data analyses: (A) Heatmap, (B) Venn diagram, and (C) tabular representation showing differential gene expression from RNA-seq analyses in ovariectomized mouse myometrium after 2 hours of the indicated treatments. The changes in expression shown in the heatmap were centered relative to the control (vehicle, V) condition and scaled relative to the maximum absolute value of the expression change for each gene (range between +1 and -1, where +1 or -1 denote the maximum or minimum values, respectively). In (B) and (C), we categorized the treatment-regulated genes with a p-value filter [p value < 0.01 (q value < 0.05)] and fold change cutoff filter (≥ 2.0 or ≤ 0.5) versus vehicle. Only genes passing both filters were included in the category. **(D)** Box plots showing the effects of the treatments on (*left*) genes uniquely regulated by E2 only (372 genes) and (*right*) genes uniquely regulated by E2+Rln (673 genes) from the Venn diagram in (B). The data are expressed relative to control, which is set to 1. Dotted red lines indicate fold change cutoff ≥ 2.0 or ≤ 0.5 (log2 ≥ 1.0 or ≤ -1.0). Boxes marked with different letters are significantly different from each other and the control (Wilcoxon rank sum test, p-value < 2.2 x 10^-16^). Genes listed below the “Upregulated” categories are representative of that category (see browser tracks in Fig. 3).

Gene ontology (GO) analysis revealed an enrichment of terms related to signaling and cell biology for the 372 E2-regulated genes that lost significant regulation upon cotreatment with Rln (Supplemental Fig. S3A). In contrast, the GO analysis revealed an enrichment of terms related to transcription and translation for the 673 E2-regulated genes that gained significant regulation upon cotreatment with Rln (Supplemental Fig. S3B). Together, these results indicate that cotreatment with Rln can alter the E2-regulated transcriptome in mouse myometrium – both by inhibiting the expression of some E2-regulated genes (e.g., *Aoc1* and *Gcat*; Fig. 3A, Supplemental Fig. S4A) and by expanding the transcriptome through increased expression of genes not regulated by E2 or Rln alone (e.g., *Sik1* and *Hapln4*; Fig. 3B, Supplemental Fig. S4B). These changes in the E2-regulated transcriptome lead to changes in downstream biological processes, as illustrated by the GO analysis (Supplemental Fig. S3).

**Figure 3.**
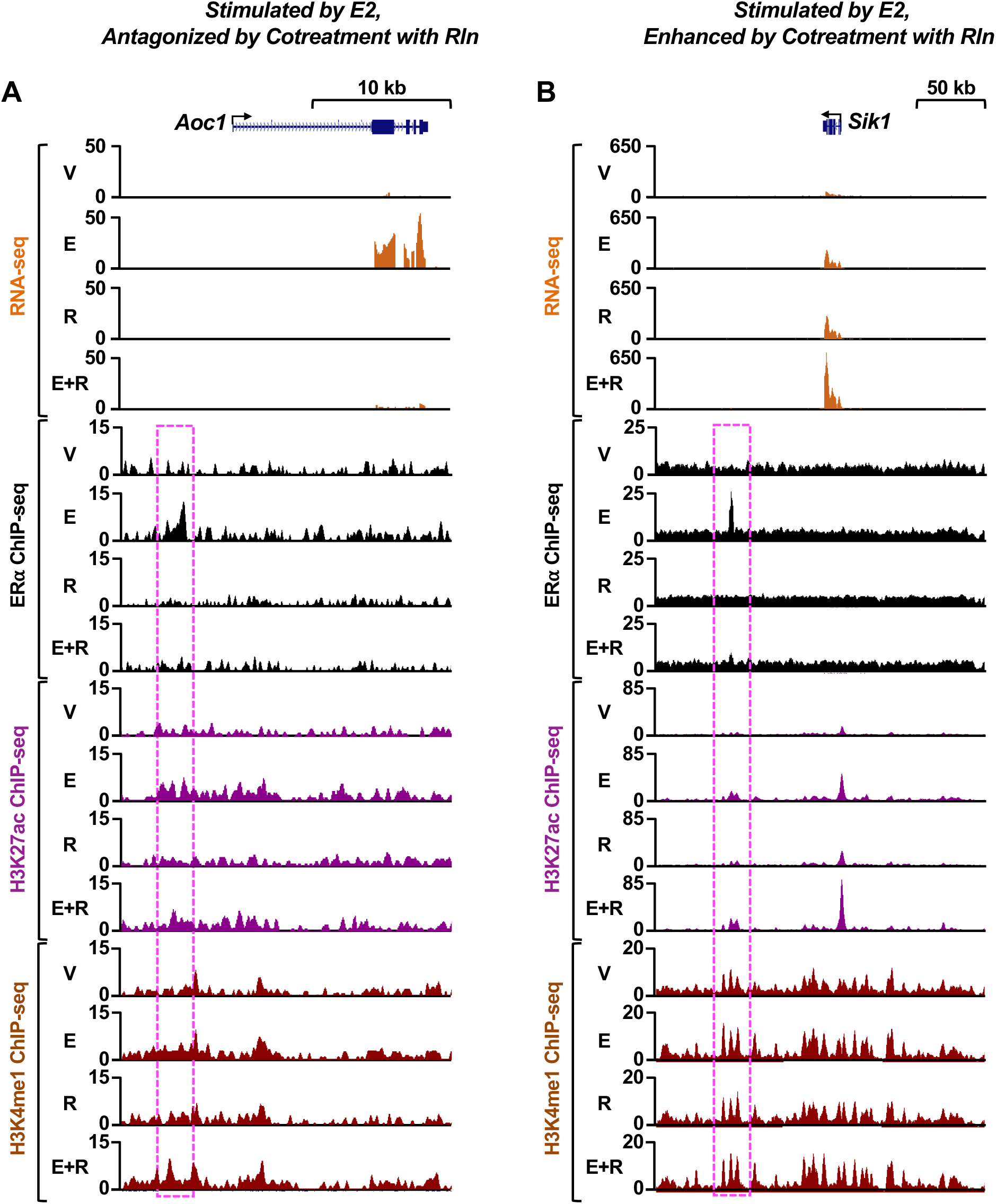
Genomic analysis of the effects of Rln treatment on genomic signaling by E2/ERα. **(A and B)** Genome browser views of RNA-seq and ERα, H3K27ac, and H3K4me1 ChIP-seq data for representative genes from ovariectomized mouse myometrium after treatment with vehicle (V), E2 (E), Rln (R), or both (E+R). Highlight (box with magenta dashed line) indicate regions of ERα enrichment from ChIP-seq. Gene schematic and scale bars are shown. The genes shown are (A) *Aoc1* (upregulated by E2 and inhibited by Rln) and (B) *Sik1* (further upregulated by E2+Rln).

### Linking the effects of Rln signaling on the ERα cistrome and the E2-regulated transcriptome in mouse myometrium

To better understand expected links between the effects of Rln signaling on the ERα cistrome and the E2-regulated transcriptome in mouse myometrium, we integrated the ChIP-seq data with the RNA-seq data to link ERα peaks depleted by Rln cotreatment with the expression of the nearest neighboring genes (Fig. 4A). We collected Rln-depleted ERα binding sites located within 50 kb of the nearest neighboring protein-coding gene promoter from the ChIP-seq data and then determined the expression of the neighboring genes using the RNA-seq data. For these analyses, we focused on genes upregulated by E2 and downregulated by cotreatment with Rln with a fold change (FC) difference (log_2_FC_E2 – log_2_FC_E+R) > 0.5. We observed a significant reduction in the E2-dependent expression of the nearest neighboring genes in response to Rln cotreatment (Fig. 4B). We then determined the ontologies for the genes associated with the Rln-depleted ERα peaks. We identified multiple GO terms related to collagen metabolism and extracellular matrix (ECM) organization (Fig. 4C; Supplemental Fig. S5A). We also identified pathways related to Rln signaling, as well as signaling via the ECM (Supplemental Fig. S5B).

**Figure 4.**
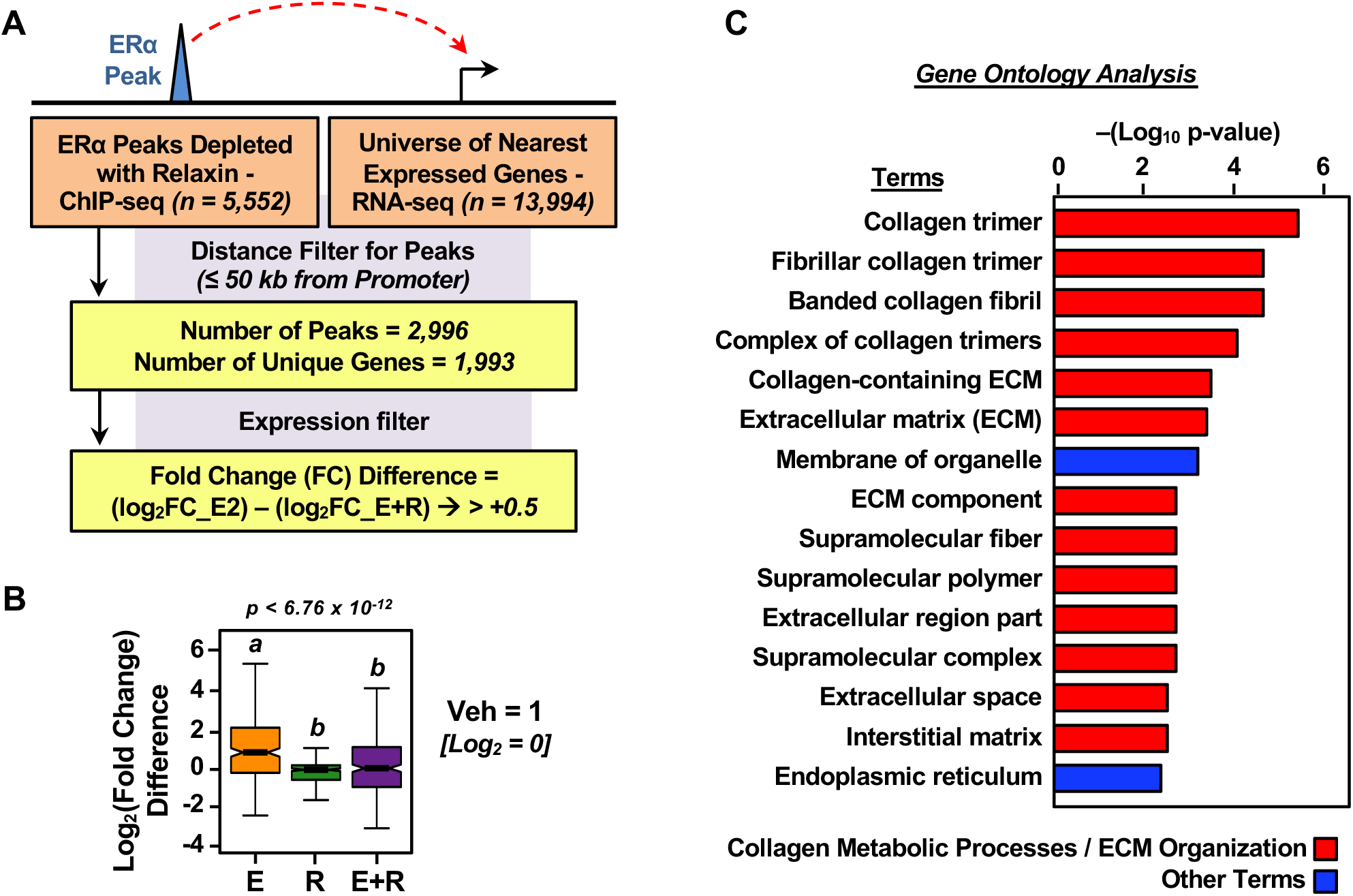
Linking the effects of Rln treatment on the ERα cistrome and E2-regulated transcriptome. **(A)** Overview of the pipeline for integrating the ChIP-seq data from Figure 1 with the RNA-seq data to link ERα peaks depleted by Rln (R) treatment with the expression of the nearest neighboring genes in mouse myometrium after treatment with vehicle (V), E2 (E), Rln (R), or both (E+R). **(B)** Box plots showing the log_2_(fold change) difference for the nearest neighboring gene expression by RNA-seq at Rln-depleted ERα binding sites. Bars marked with different letters are significantly different (Wilcoxon rank sum test, p < 0.001). **(C)** Gene ontologies for the genes associated with Rln-depleted ERα peaks were evaluated using WEB-based GEne SeT AnaLysis Toolkit (WebGestalt) (59). Multiple gene ontology terms related to collagen metabolism and extracellular matrix (ECM) organization were identified.

Collectively, the ChIP-seq and RNA-seq results demonstrate that the Rln-dependent changes that we observed in the E2-dependent ERα cistrome are reflected in the gene expression outcomes and are connected to biological outcomes related to Rln signaling. This is evident for genes encoding amine oxidase copper containing 1 (*Aoc1*) and glycine C-acetyltransferase (*Gcat*), both of which exhibit reduced E2-dependent ERα binding at a nearby enhancer, as well reduced expression, upon cotreatment with Rln (Fig. 3A, Supplemental Fig. S4A). However, Rln-mediated modulation of the E2-dependent ERα cistrome (depletion of >5,000 ERα binding sites; Fig. 1D) was more extensive than Rln-mediated modulation of the E2-dependent transcriptome (reduced expression of ∼370 genes; Fig. 2, A and B). In this regard, classical E2/ERα-regulated genes, such as *Pgr*, *Nrip1*, and *Igf1* exhibited a loss of E2-dependent ERα binding at a nearby enhancer upon cotreatment with Rln without showing an observable effect on expression (Fig. 5; Supplemental Fig. S6). Importantly, the inhibitory effects of Rln on the E2-dependent binding of ERα at enhancers was abrogated by genetic knockout of *Rxfp1* (52), the gene encoding the Rln receptor (Supplemental Fig. S7), reinforcing the role of Rln acting through its receptor for the modulation of E2/ERα signaling, as well as the resulting downstream transcriptional effects and biological outcomes.

**Figure 5.**
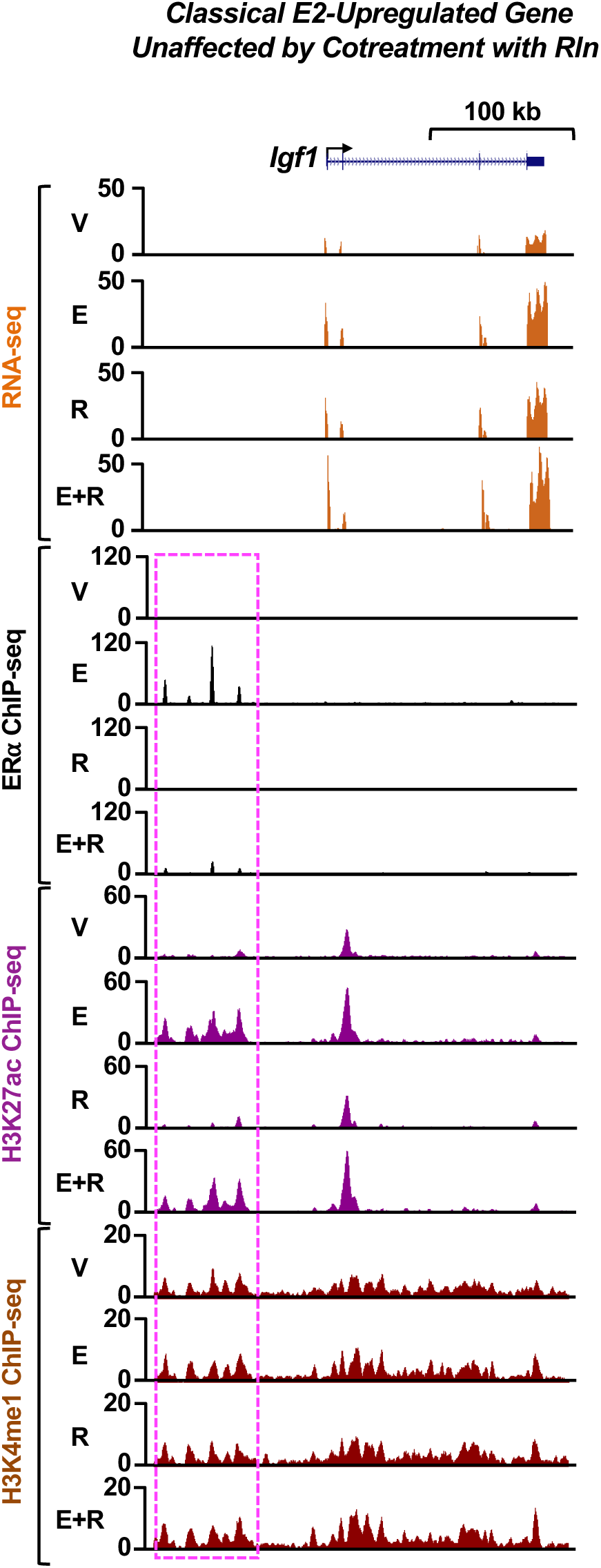
Genomic analysis of the effects of Rln treatment on genomic signaling by E2/ERα at a classical estrogen-regulated gene. Genome browser views of RNA-seq and ERα, H3K27ac, and H3K4me1 ChIP-seq data for the classical E2-regulated gene *Igf1* from ovariectomized mouse myometrium after treatment with vehicle (V), E2 (E), Rln (R), or both (E+R). Highlight (box with magenta dashed line) indicates regions of ERα enrichment from ChIP-seq. Gene schematic and scale bars are shown.

### The Rln receptor and ERα phosphorylation mediate the inhibitory effects of Rln on estrogen signaling in myometrial tissue and cells

We considered the possibility that receptor-mediated Rln signaling might ultimately connect to the E2 signaling pathway through phosphorylation of ERα, which has been shown to regulate ERα-dependent gene expression (43,44,75). ERα is phosphorylated on multiple amino acid residues, including three prominent sites in its unstructured amino-terminal region: Ser118, Ser104/106, and Ser167 (43). Ser118 and Ser104/106 are the predominant E2-dependent phosphorylation sites, with phosphorylation of Ser118 thought to occur in response to activation of mitogen-activated protein kinases (MAPKs) (43). The latter was determined in breast cancer cells, but has not be explored in detail in myometrium. We observed a dramatic increase in Ser118 phosphorylation in response to E2 treatment in wild-type mouse myometrium, as indicated by Western blotting (Fig. 6A). We also observed a more modest enhancement of Ser104/106 phosphorylation upon E2 treatment, but we did not observe a significant change in Ser167 phosphorylation (Fig. 6A; Supplemental Fig. S8). Cotreatment with Rln abrogated the E2-mediated increase in Ser118 phosphorylation and enhanced the E2-mediated increase in Ser104/106 phosphorylation (Fig. 6A; Supplemental Fig. S8). The effects of Rln were dependent on the Rln receptor, as demonstrated in studies using *Rxfp1^-/-^* mice (Fig. 6A).

**Figure 6.**
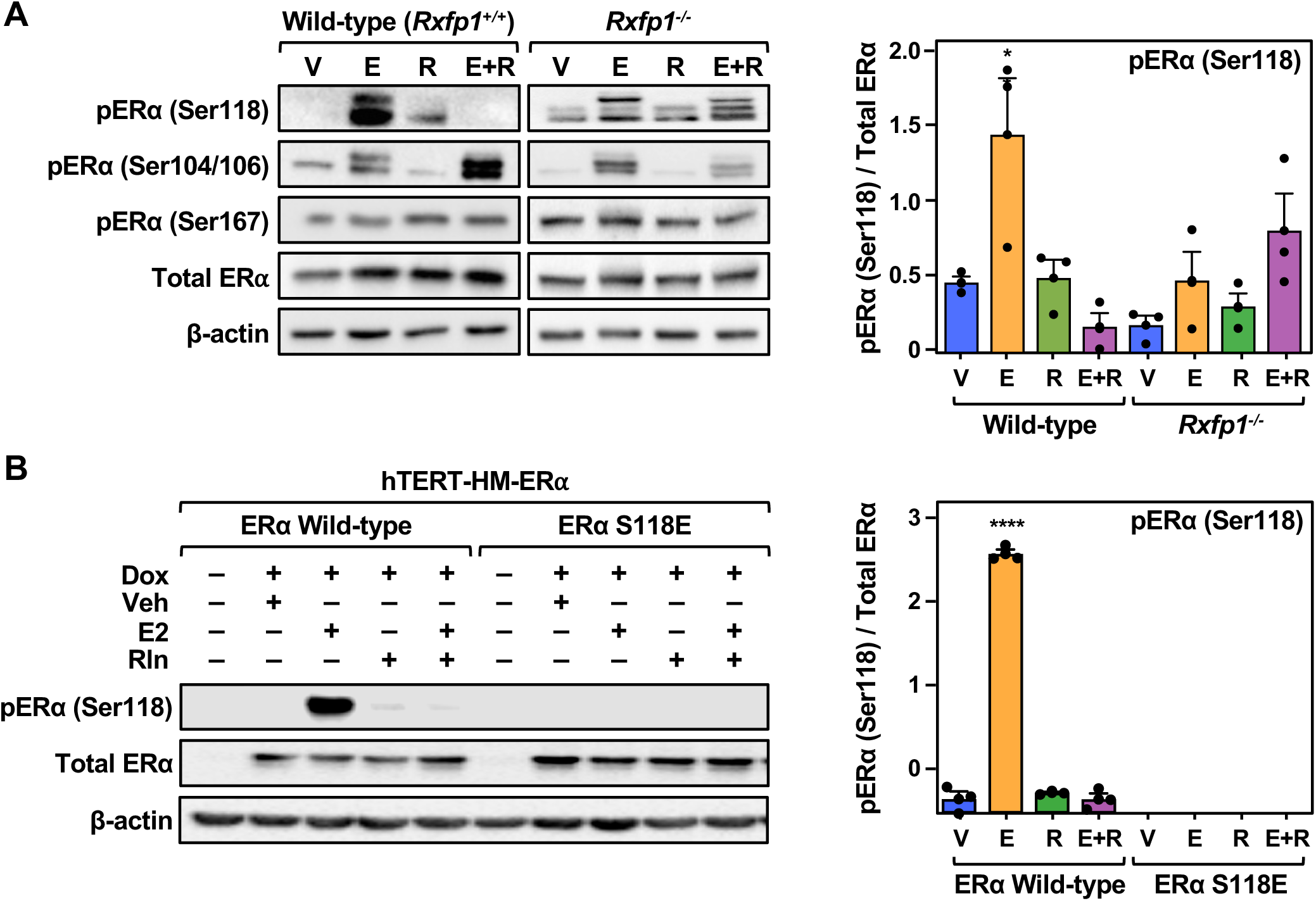
The effects of Rln on estrogen signaling in myometrial cells requires inhibition of ERα phosphorylation at serine 118. **(A)** Western blotting *(left)* and densitometric quantification *(right)* of ERα phosphorylation levels at Ser118, Ser104/106, and Ser167 in myometrium from ovariectomized wild-type and Rln receptor knockout (*Rxfp1^-/-^*) mice treated for 1 hour with the treatments indicated: vehicle (V), E2 (E), Rln (R), or both (E+R). β-actin serves as a loading control. Each bar represents the mean + SEM, n = 3, 5 mice/treatment). The asterisk indicates significant difference from the corresponding control (V) (One-way ANOVA, * p < 0.05). **(B)** Western blotting *(left)* and densitometric quantification *(right)* of ERα phosphorylation levels at Ser118 in Dox-inducible hTERT-HM-ERα cells with wild-type or phosphomimetic S118E mutant ERα subjected to the treatments indicated for 30 minutes: vehicle (V), E2 (E), Rln (R), or both (E+R). β-actin serves as a loading control. Each bar represents the mean + SEM, n = 3. The asterisks indicate significant difference from the corresponding control (V) (One-way ANOVA, **** p < 0.0001).

To determine if similar effects of Rln cotreatment on ERα Ser118 phosphorylation occur in human myometrial cells, we used telomerase-immortalized human myometrial (hTERT-HM) cells (53), which we engineered with doxycycline (Dox)-inducible expression of ERα (wild-type, WT) and ERα mutants (S118A and S118E) by lentivirus-mediated transduction. These cells show similarities to mouse myometrial cells in their responses to E2 and Rln based on gene expression (Supplemental Fig. S9) and GO analysis (Supplemental Fig. S10). Moreover, hTERT-HM cells expressing the ERα S118E mutant exhibited increased basal (i.e., without E2) expression of an E2-regulated gene set and a loss of Rln-mediated inhibition of the same gene set (Supplemental Fig. S11).

As with the mouse myometrium, we observed a dramatic increase in Ser118 phosphorylation in response to E2 treatment in the presence of wild-type ERα (Fig. 6B). The effect of E2 on S118 phosphorylation was blocked by cotreatment with Rln, as well as mutation of S118 to glutamate (S118E) or alanine (S118A) (Fig. 6B; Supplemental Fig. S12A).

Importantly, although S118E blocks the epitope recognized by the phosphorylated ERα Ser118 antibody, this mutant served as a useful phosphomimetic in the studies described below, in contrast to the S118A mutant, due to the negatively charged glutamate. Importantly, an inhibitor of the mitogen-activated protein kinase kinase (MEK), Trametinib (GSK1120212), which blocks signaling through the MAPK/ERK pathway (51), had no observable effect on E2-induced ERα S118 phosphorylation or its inhibition by cotreatment with Rln (Supplemental Fig. S12B). Together, these studies link ERα S118 phosphorylation and global gene expression outcomes, but indicate the phosphorylation event is not mediated by a MAPK/ERK pathway in myometrial cells.

### The inhibitory effects of Rln on ERα Ser118 phosphorylation blocks the E2-induced contraction of myometrial cells

To explore the downstream biology of the inhibitory effects of Rln on ERα Ser118 phosphorylation, we used a collagen gel contraction assay with hTERT-HM-ERα cells expressing wild-type or S118E mutant ERα. This assay measures the ability of the myometrial cells to contract *in vitro*. Briefly, hTERT-HM-ERα cells (WT, S118A and S118E) with Dox-induced ERα expression were plated in collagen with subsequent incubation for 1 hour at 37°C to allow gelling. The cells were then treated in fresh medium with 100 nM E2, 0.2 μg/mL Rln, or a combination of E+R. After 20 hours, the gel matrices were gently detached from the wells and the areas of the gel matrices were measured periodically for up to 48 hr. Using this assay, we observed a potent effect of E2 on the induction of contraction in cells expressing wild-type ERα [Percent contraction: 24 ± 0.8 (V) versus 55 ± 1.5 (E2)] , which was completely blocked by cotreatment with Rln [Percent contraction: 55 ± 1.5 (E2) versus 24 ± 0.5 (E+R)] (Fig. 7). We observed a similar stimulatory effect of E2 in cells expressing ERα S118E, which was unaffected by cotreatment with Rln [Percent contraction: 58 ± 1.3 (E2) versus 58 ± 1.0 (E+R)] (Fig. 7). In contrast, no contraction was observed in cells expressing ERα S118A (Supplemental Fig. S12C). These results indicate that E2-induced contraction of myometrial cells requires phosphorylation of ERα at Ser118 and that Rln blocks this effect by inhibiting Ser118 phosphorylation.

**Figure 7.**
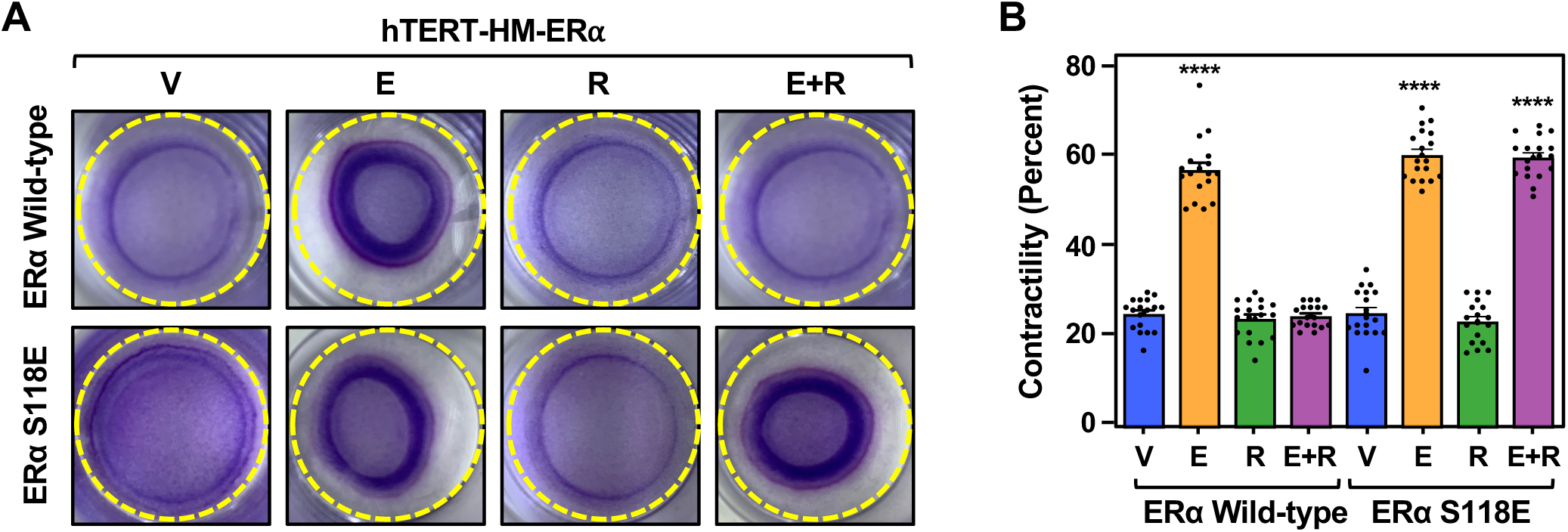
The effects of Rln on an estrogen-dependent biological outcome in myometrial cells requires inhibition of ERα phosphorylation at serine 118. **(A and B)** Representative images (A) and quantification (B) of collagen gel contraction assays conducted in hTERT-HM-ERα cells with wild-type or S118E mutant ERα. The cells were plated in collagen with subsequent incubation for 1 hour at 37°C to allow gelling. The cells were then treated in fresh medium with vehicle (V), E2 (E), Rln (R), or both (E+R). After 20 hours, the gel matrices were gently detached from the wells and the areas of the gel matrices were measured periodically for up to 48 hr. The area of each gel was measured after 48 hours and plotted as percent contraction compared to the area of untreated well at the beginning of experiment. Each bar represents the mean + SEM, n = 3, 6 wells/treatment. The asterisks indicate significant difference from the corresponding control (V) (One-way ANOVA, **** p < 0.0001).

### DUSP1 and DUSP5 are required for Rln-mediated inhibition of E2-induced ERα Ser118 phosphorylation and contraction in myometrial cells

We sought to connect the effects of Rln treatment to a molecular mechanism that could inhibit E2-induced phosphorylation of ERα at Ser118. We considered the possibility that a Rln-regulated phosphatase might act to dephosphorylate Ser118, with nuclear dual-specificity phosphatases (DUSPs) being good candidates. DUSPs dephosphorylate many key signaling molecules, thereby regulating the magnitude and duration of phosphorylation-dependent events (76,77). In this regard, a previous study demonstrated that E2-induced phosphorylation and activation of ERα was inhibited by catalytically active DUSP22, but not catalytically inactive mutants, in breast cancer cells (78). In our studies in myometrial cells, we focused on two nuclear DUSPs, DUSP1 and DUSP5 (2,79,80), which are expressed in hTERT-HM-ERα cells and whose expression is unaffected by the treatments used in this study (E2, Rln, E+R) (Supplemental Fig. S13A).

We used siRNA-mediated knockdown to deplete DUSP1, DUSP5, or both in hTERT-HM-ERα cells (Supplemental Fig. S13, B and C), then examined the effect on E2-induced phosphorylation of ERα at Ser118, as well as its reversal by cotreatment with Rln. Depletion of DUSP1, DUSP5, or both in combination alone had little effect on E2-induced Ser118 phosphorylation, but completely blocked the inhibition of Ser118 phosphorylation by Rln (Fig. 8A). In the contraction assay, treatment of hTERT-HM cells expressing wild-type ERα and control siRNA (siControl) again showed a robust induction of contraction [Percent contraction: 24 ± 0.7 (V) versus 54 ± 0.4 (E2)] and inhibition by cotreatment with Rln [Percent contraction: 54 ± 0.4 (E2) versus 23 ± 0.6 (E+R)] (Fig. 8, B and C). Depletion of DUSP1 alone had little effect on E2-induced contraction or inhibition by cotreatment with Rln [Percent contraction: 23 ± 0.5 (V), 55 ± 0.6 (E2), 27 ± 0.1 (E+R)], whereas depletion of DUSP5 alone enhanced E2-induced contraction and partially reversed inhibition by cotreatment Rln [Percent contraction: 30 ± 0.3 (V), 64 ± 0.4 (E2), 40 ± 0.9 (E+R)] (Fig. 8, B and C). Depletion of both DUSP1 and DUSP5 enhanced E2-induced contraction and reversed inhibition by cotreatment with Rln [Percent contraction: 28 ± 0.4 (V), 68 ± 0.3 (E2), 55 ± 0.5 (E+R)] (Fig. 8, B and C). Depletion of DUSP1, DUSP5, or both in combination also modestly enhanced the effects of Rln treatment alone (Fig. 8, B and C). These results implicate both DUSP1 and DUSP5 as key mediators of the inhibitory effects of Rln on ERα, with perhaps a more prominent role for DUSP5.

**Figure 8.**
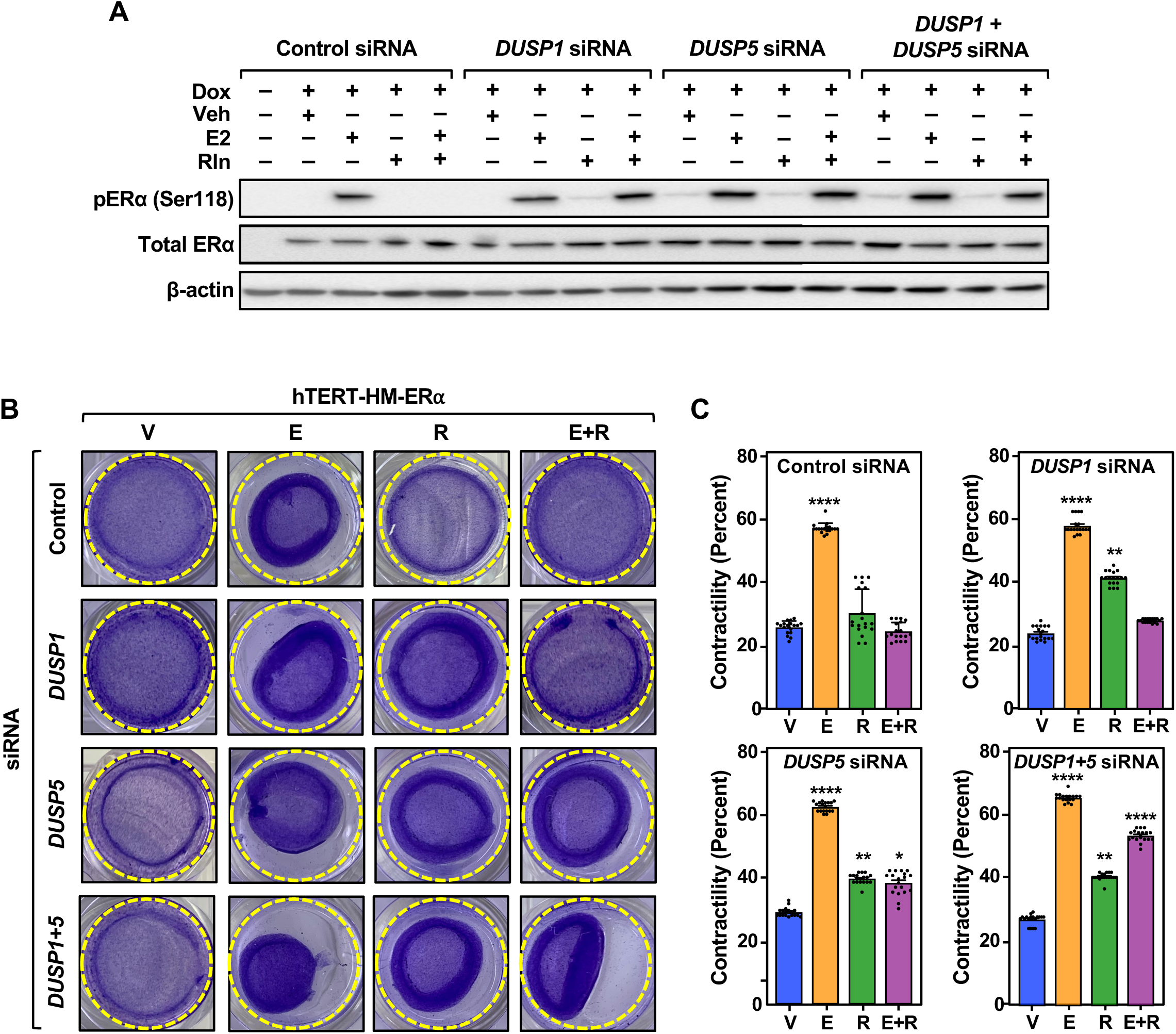
DUSP1 and DUSP5 are required for Rln-mediated inhibition of E2-induced ERα Ser118 phosphorylation and contraction in myometrial cells. **(A)** Western blotting of phospho ERα Ser118 and total ERα in hTERT-HM-ERα cells with siRNA-mediated knockdown of *DUSP1* mRNA, *DUSP5* mRNA or both. The cells were subjected to the treatments indicated for 30 minutes: vehicle (V), E2 (E), Rln (R), or both (E+R). β-actin serves as a loading control. **(B)** Representative images from collagen gel contraction assays conducted in hTERT-HM-ERα cells with siRNA-mediated knockdown of *DUSP1* mRNA, *DUSP5* mRNA or both. The cells were subjected to the treatments indicated as described in the Methods: vehicle (V), E2 (E), Rln (R), or both (E+R). **(C)** Quantification of the collagen gel contraction assays shown in (B). The area of each gel was measured after 48 hours and plotted as percent contraction compared to the area of untreated well at the beginning of experiment. Each bar represents the mean + SEM, n = 3, 6 wells/treatment. The asterisks indicate significant difference from the corresponding control (V) (One-way ANOVA, * p < 0.05, ** p < 0.01, **** p < 0.0001).

## Discussion

In this study, we explored the molecular mechanisms by which Rln/RXFP1 signaling can promote or oppose the actions of E2/ERα in myometrial cells and tissue, which is a poorly studied aspect of reproductive tract biology. Our results indicate that Rln attenuates the genomic actions and biological effects of estrogen in the myometrium by reducing ERα S118 phosphorylation. Interestingly, we observed a potent inhibitory effect of Rln/RXFP1 on ERα binding across the genome. The reduction in ERα binding was associated with changes in the hormone-regulated transcriptome, including a decrease in the E2-dependent expression of some neighboring genes, but not selected classical E2-regulated genes (Fig. 9). Cotreatment with E2 and Rln also promoted an expansion of the transcriptome by further enhancing the expression of E2-regulated genes beyond that observed with E2 or Rln alone (Fig. 9). The effects of Rln cotreatment were manifested in a reduction in the E2-dependent phosphorylation of ERα on Ser 118, as well the E2-dependent contraction of myometrial cells. Importantly, we observed a requirement for DUSP1 and DUSP5 for the inhibitory effects of Rln cotreatment on estrogen signaling. Collectively, our results identify a pathway that integrates Rln/RXFP1 and E2/ERα signaling, resulting in a convergence of membrane and nuclear signaling pathways to control genomic and biological outcomes (Fig. 9).

**Figure 9.**
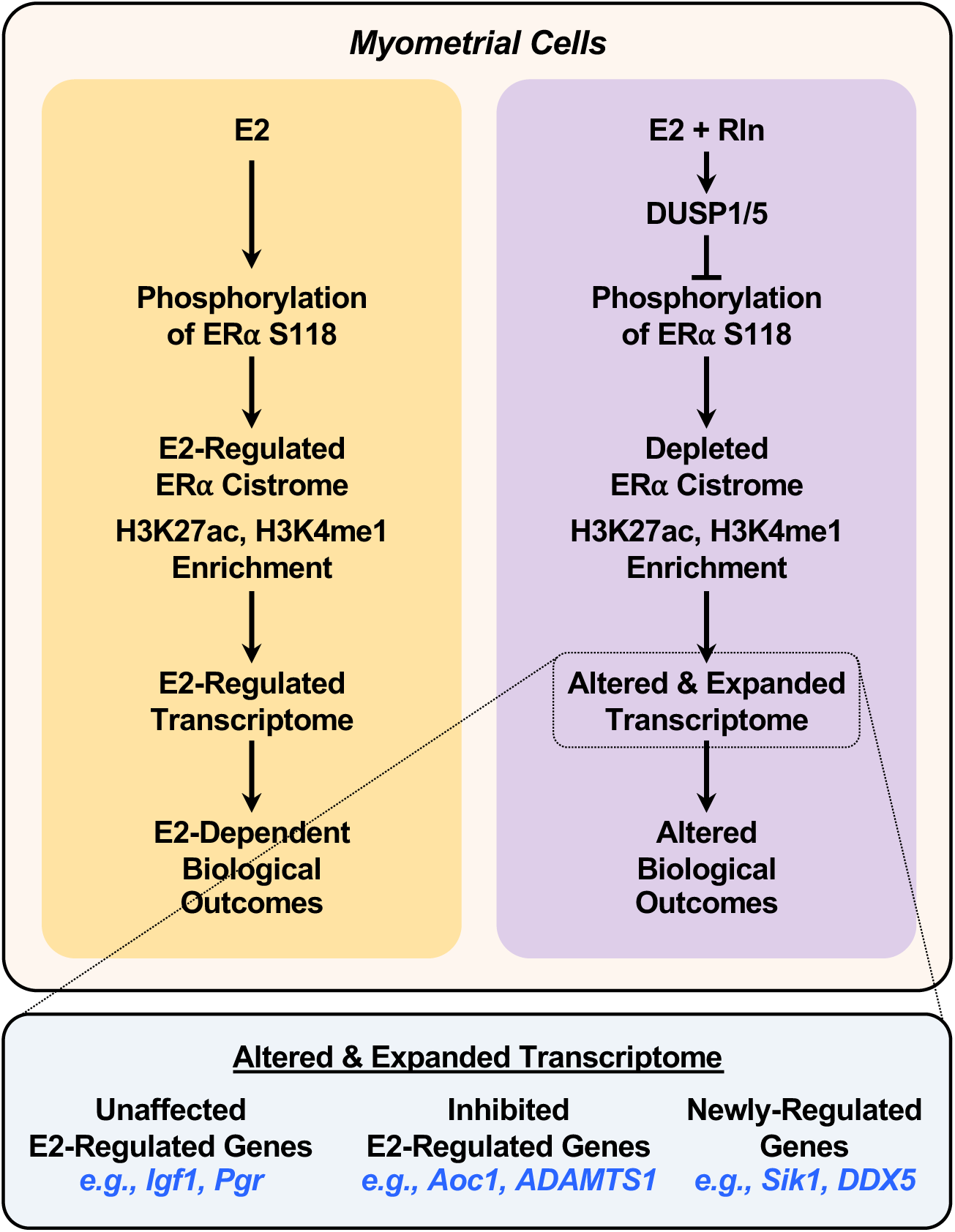
Summary of the observations regarding the effects of Rln on ERα-dependent gene regulation in myometrial cells. Additional components (e.g., DUSPs 1 and 5) are involved in E2/ERα signaling upon cotreatment with Rln. Cotreatment with Rln alters and expands the E2-regulated transcriptome, as indicated in the bottom panel. See the text for details.

### Molecular mechanisms underlying the unique regulatory effects of Rln on estrogen signaling in the myometrium

Previous studies have characterized both stimulatory and inhibitory crosstalk between Rln and E2 signaling on biological outcomes in various reproductive tissues. For example, many of the effects of Rln on reproductive tissues, such as cell proliferation, ECM remodeling, and angiogenic actions, require or are augmented by E2/ERα signaling (16,26,27). In contrast, Rln promotes smooth muscle quiescence in the myometrium, while E2/ERα promotes contraction (2,28–30). Our studies suggest a potential mechanism for the opposing actions, namely the control of E2-dependent gene regulation by altering ERα phosphorylation status and binding across the genome. In this regard, a phosphomimetic ERα mutant S118E enhanced basal ERα-dependent gene expression and conferred resistance to the inhibitory effects of Rln. We also found that DUSP1 and DUSP5 mediate the reduction in E2-dependent phosphorylation of ERα on Ser118 with Rln cotreatment (Fig. 9). Ultimately, this leads to reduced contraction of myometrial cells. Interestingly, another DUSP, DUSP22, was previously shown to regulate ERα-mediated signaling via direct dephosphorylation of Ser118 (78). Previous studies in breast cancer cells showed that phosphorylation of ERα on Ser118 occurs predominantly in response to activation of MAPKs (43). In contrast, our studies herein using a MEK inhibitor did not recapitulate these results in myometrial cells, leaving open the question of the kinase that phosphorylates ERα in these cells. Nonetheless, as we show herein, inhibition of E2-dependent phosphorylation of ERα by cotreatment with Rln can be reversed by the actions of DUSP1 and DUSP5 (Fig. 9).

### Limitations of the study and future directions

Some of the best characterized effects of Rln are during pregnancy, where it acts to relax the uterine muscle and soften the pelvic ligaments to facilitate parturition (15–17). In our ovariectomized, non-pregnant mouse model, we were unable to investigate the functional interplay between Rln/RXFP1 and E2/ERα signaling during pregnancy and parturition. This will be an interesting direction to pursue in future studies. Regarding the molecular mechanisms by which DUSP1 and DUSP5 inhibit the phosphorylation of ERα on Ser118, our results and previous studies (78) suggest that this might occur through the direct actions of the DUSPs on phosphorylated ERα, but we have not shown this directly in biochemical assays. In addition, as noted above, we have identified the enzyme that mediates ERα phosphorylation. These will be additional interesting questions to address in future studies.

## Acknowledgments

The authors would like to acknowledge and thank the following: Dr. Carole Mendelson for the hTERT-HM cells; Dr. Alexander Agoulnik for the *Rxfp1^-/-^* mice; Asha Varghese, Cosette Taggart, and Carla De Cassia Villela for assistance with mouse genotyping; ShanmugaPriyaa Madhukaran for assistance with ovariectomy surgery and members of the Kraus and Mahendroo laboratories for input and feedback on this project. We would also like to acknowledge the UT Southwestern Next Generation Sequencing Core, under the direction of Ralf Kittler.

## Dedication

The authors would like to dedicate this manuscript to the memory of Dr. Carole Mendelson, a beloved colleague and friend: https://www.utsouthwestern.edu/education/medical-school/departments/green-center/assets/in-memoriam-carole-mendelson.pdf

## Author Contributions

ST - Conceptualization, Methodology, Validation, Formal analysis, Investigation, Writing – original draft preparation, Visualization.

AN - Software, Formal analysis, Data Curation, Writing – original draft preparation, Visualization.

SPC - Formal analysis, Investigation.

TN - Formal analysis, Writing – review and editing, Visualization.

CVC - Methodology, Writing – review and editing, Supervision.

MM - Conceptualization, Methodology, Writing – review and editing, Visualization, Supervision, Project administration, Funding acquisition.

WLK - Conceptualization, Methodology, Writing – review and editing, Visualization, Supervision, Project administration, Funding acquisition.

## Data Availability

The genomic datasets generated for this study can be accessed from the NCBI’s Gene Expression Omnibus (GEO) repository (www.ncbi.nlm.nih.gov/geo/) using superseries accession number GSE244843. All other data generated in this study are available within the article and its supplementary data files. The pipelines and scripts used for the computational analyses are available on request.

## SUPPLEMENTARY DATA FILES

This paper contains Supplementary Figures S1-S13.

## Abbreviations

E2: estradiol
Rln: relaxin
RXFP1: relaxin family peptide receptor 1 (relaxin receptor)
DUSP: dual-specificity phosphatase
ChIP: chromatin immunoprecipitation
RT-qPCR: reverse transcription-quantitative PCR.

