## Supplemental Figures 1-13 for "Relaxin Modulates the Genomic Actions and Biological Effects of Estrogen in the Myometrium"

**Tripathy *et al.*, 2024**

This file contains:
- Supplemental Figures S1 through S13
- Supplemental Figure Legends S1 through S13

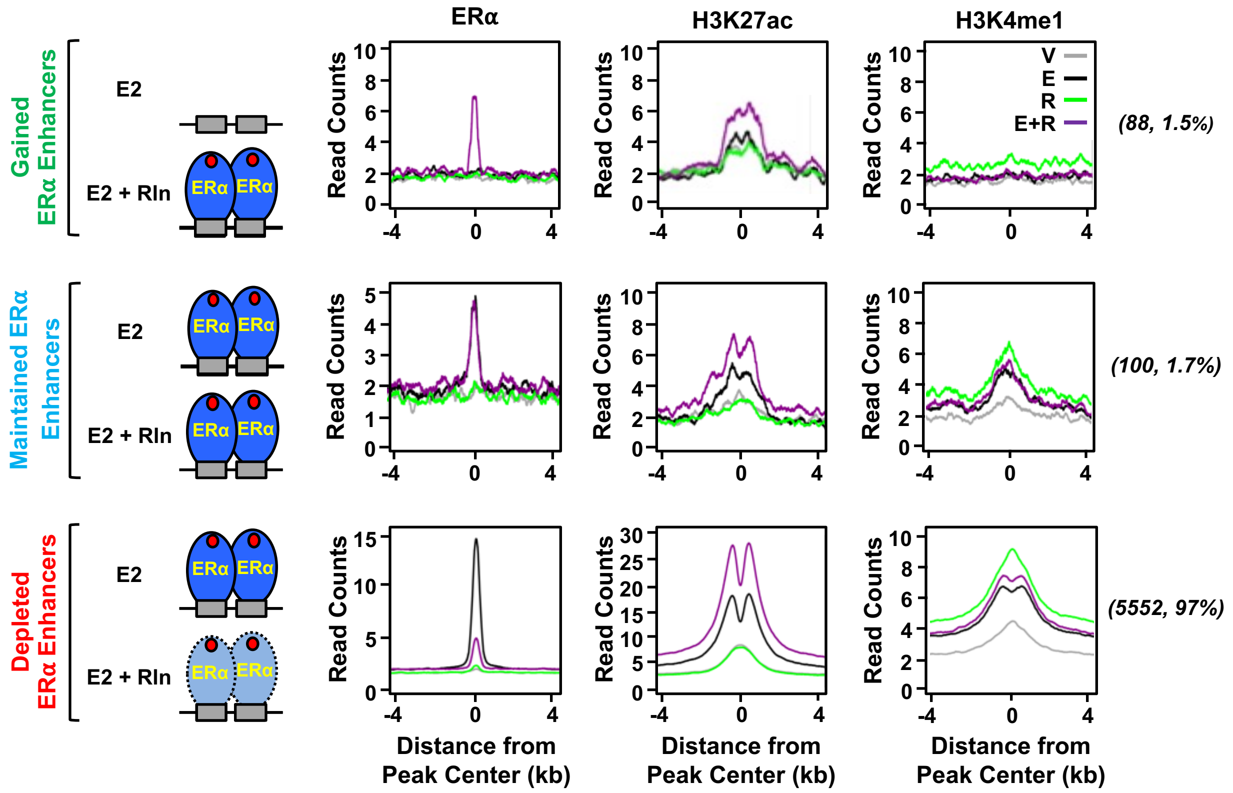


**Supplemental Figure S1. Metaplots of ERα, H3K27ac, H3K4me1 ChIP-seq data from mouse myometrium after treatment with E2 and Rln.**
Metaplot analysis showing relationships between H3K27ac and H3K4me1 enrichment for the uniquely bound ERα sites in myometrium isolated from ovariectomized mice treated with E2 (E), Rln (R) or both (E+R).

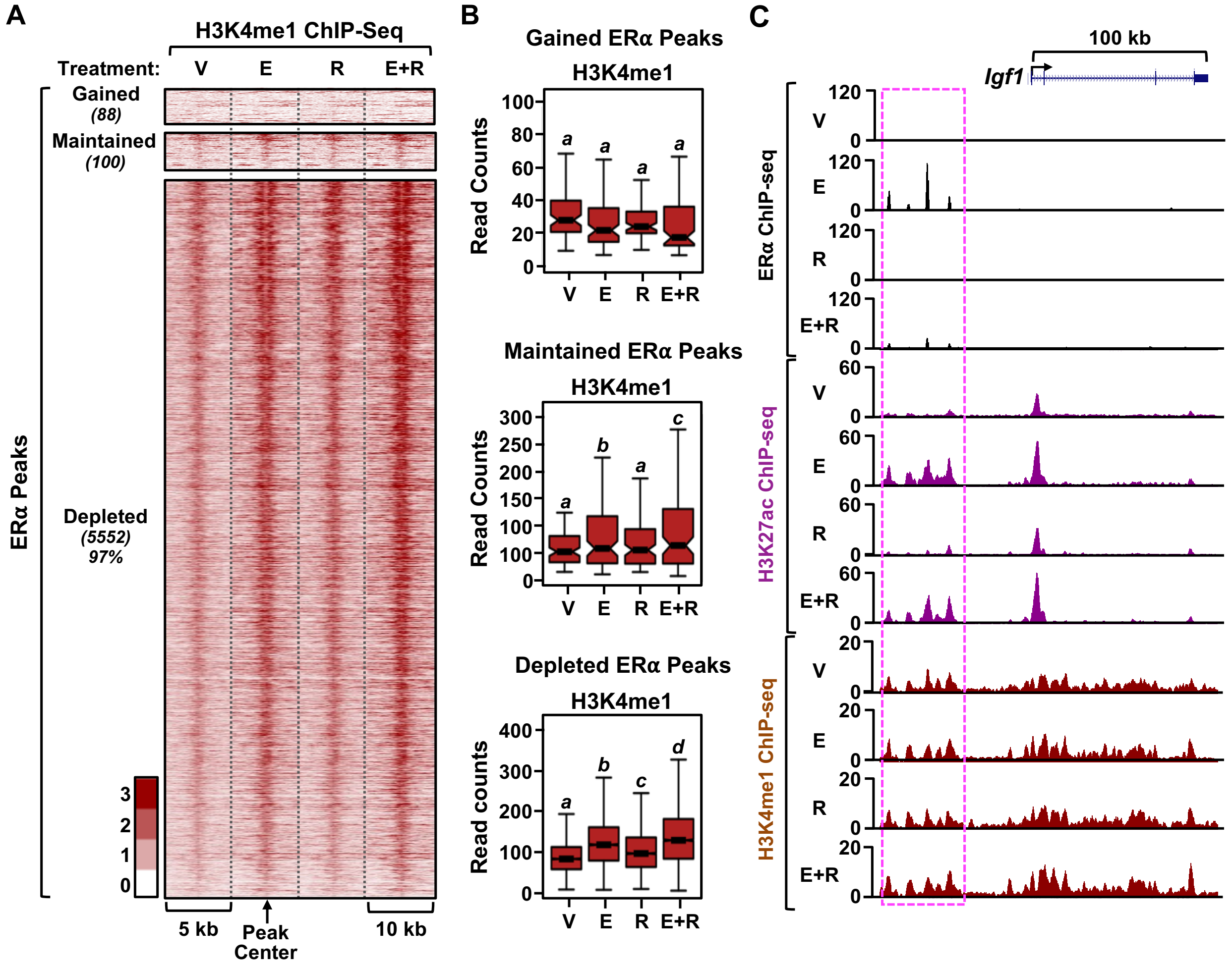


**Supplemental Figure S2. The levels of H3K4me1 are unchanged in mouse myometrium after treatment with E2 and Rln.**

**(A)** Heatmaps of H3K4me1 ChIP-seq data at ERα binding sites in myometrium obtained from ovariectomized mice treated with vehicle (V), E2 (E), Rln (R), or both (E+R). The changes shown in the heatmap were centered relative to the control (vehicle, V) condition.

**(B)** Box plots of normalized read counts for H3K4me1 ChIP-seq data at three distinct sets of ERα binding sites (gained, maintained, lost). Bars marked with different letters are significantly different (Wilcoxon rank sum test, $p < 0.001$).

**(C)** Genome browser views of ERα, H3K27ac, and H3K4me1 ChIP-seq data for the *Igf1* gene from mouse myometrium after treatment with vehicle (V), E2 (E), Rln (R), or both (E+R). The highlight (box with magenta dashed line) indicate a region of ERα enrichment from ChIP-seq. Gene schematic and scale bar are shown.

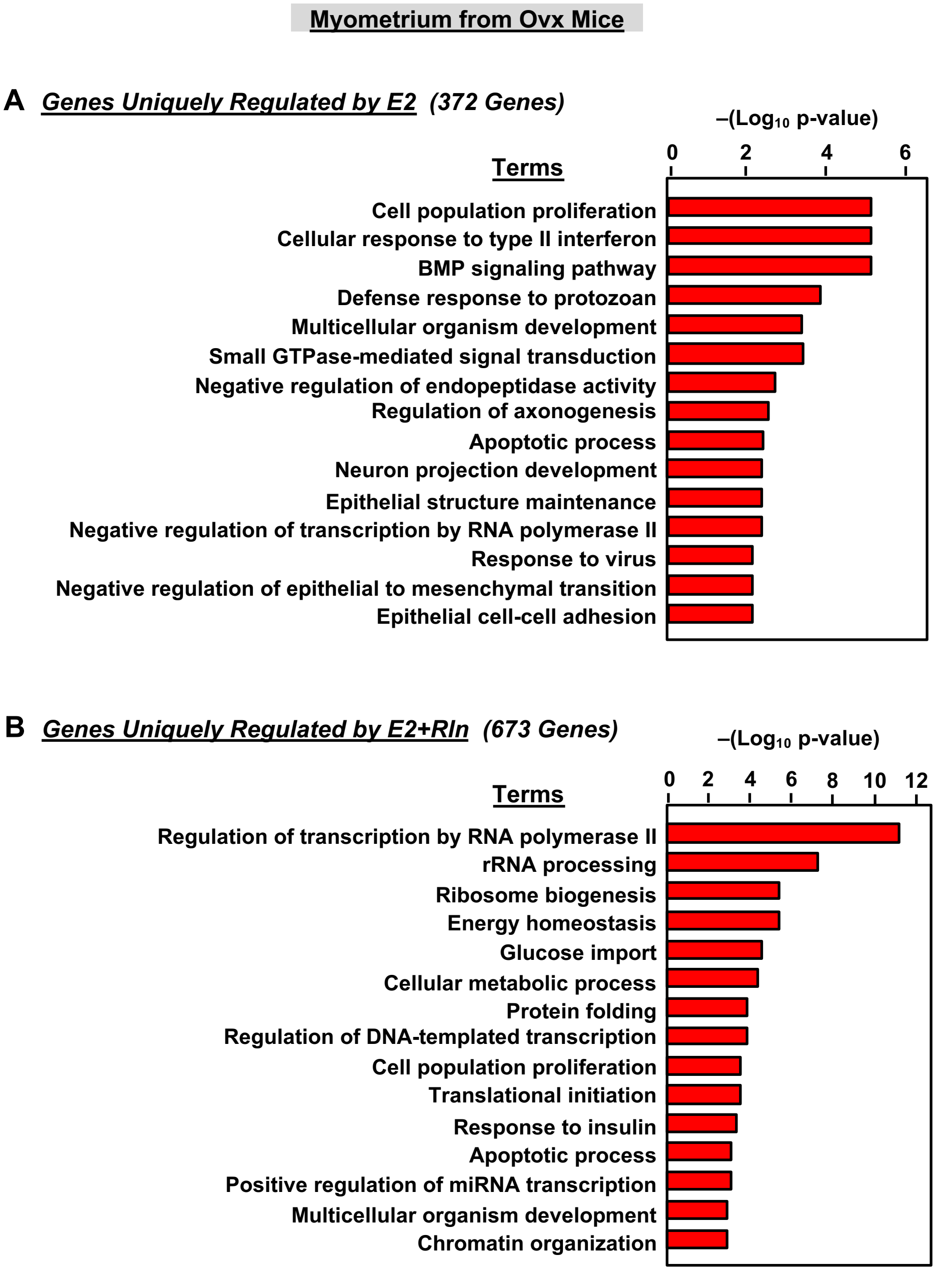


**Supplemental Figure S3. Gene ontologies for E2- and Rln-regulated genes in myometrium from ovariectomized mice.**
GO analyses of uniquely regulated genes from the Venn diagram in Figure 2B. The gene sets were evaluated using WEB-based Gene SeT AnaLysis Toolkit (WebGestalt) (59).
**(A)** GO analysis for genes uniquely regulated by E2 (372 genes).
**(B)** GO analysis for genes uniquely regulated by E2+Rln (673 genes).

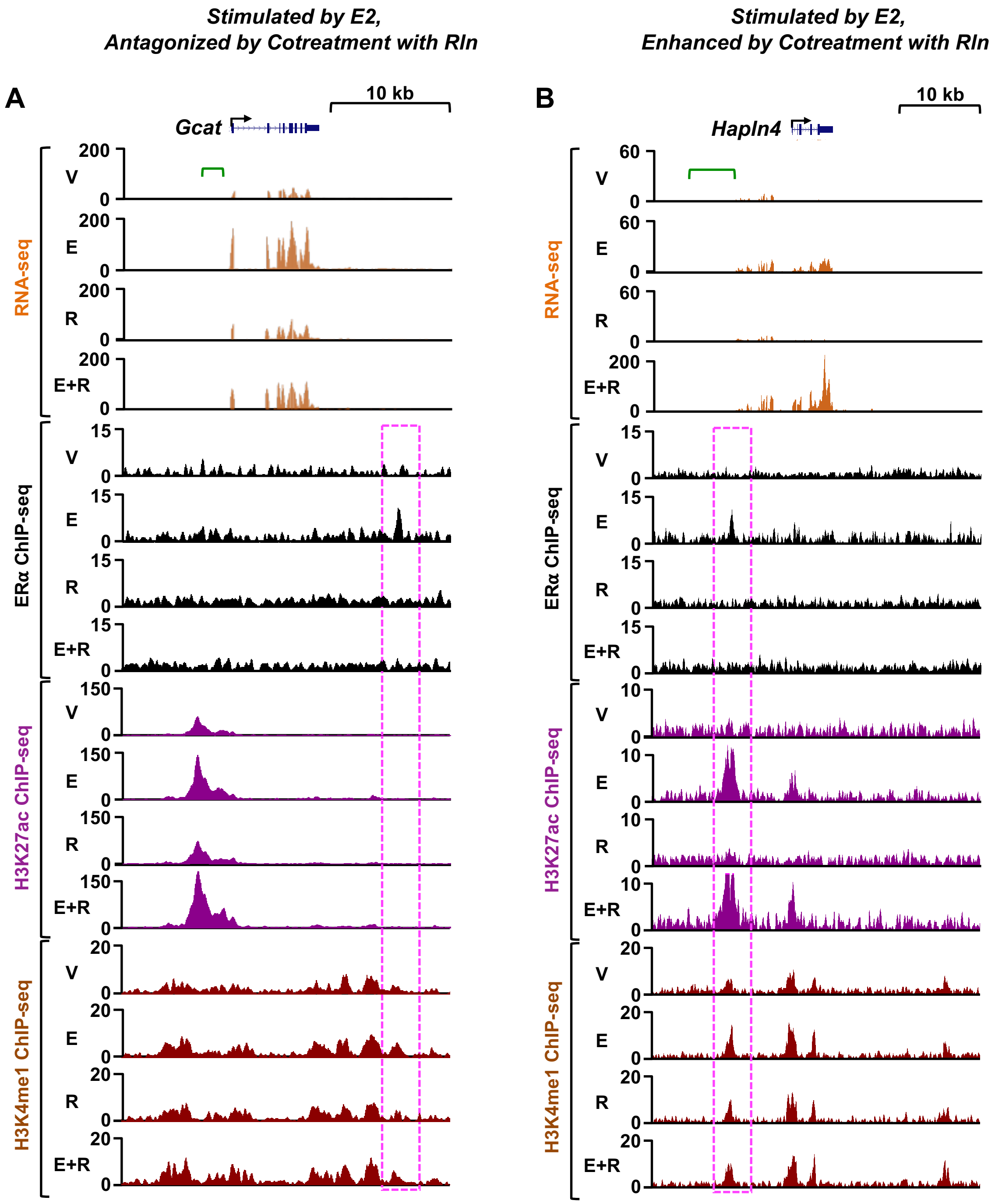


**Supplemental Figure S4. Genomic analysis of the effects of Rln treatment on genomic signaling by E2/ERα.**

**(A and B)** Genome browser views of RNA-seq and ERα, H3K27, and H3K4me1 ChIP-seq data for representative genes from ovariectomized mouse myometrium after treatment with vehicle (V), E2 (E), Rln (R), or both (E+R). Highlight (box with magenta dashed line) indicate regions of ERα enrichment from ChIP-seq. Gene schematic and scale bars are shown. Green brackets indicate regions outside of the gene of interest that have been removed from the RNA-seq browser tracks due to excessive signal that dwarfed the signal of interest. The genes shown are (A) *Gcat* (upregulated by E2 and inhibited by Rln) and (B) *Hapln4* (further upregulated by E2+Rln).

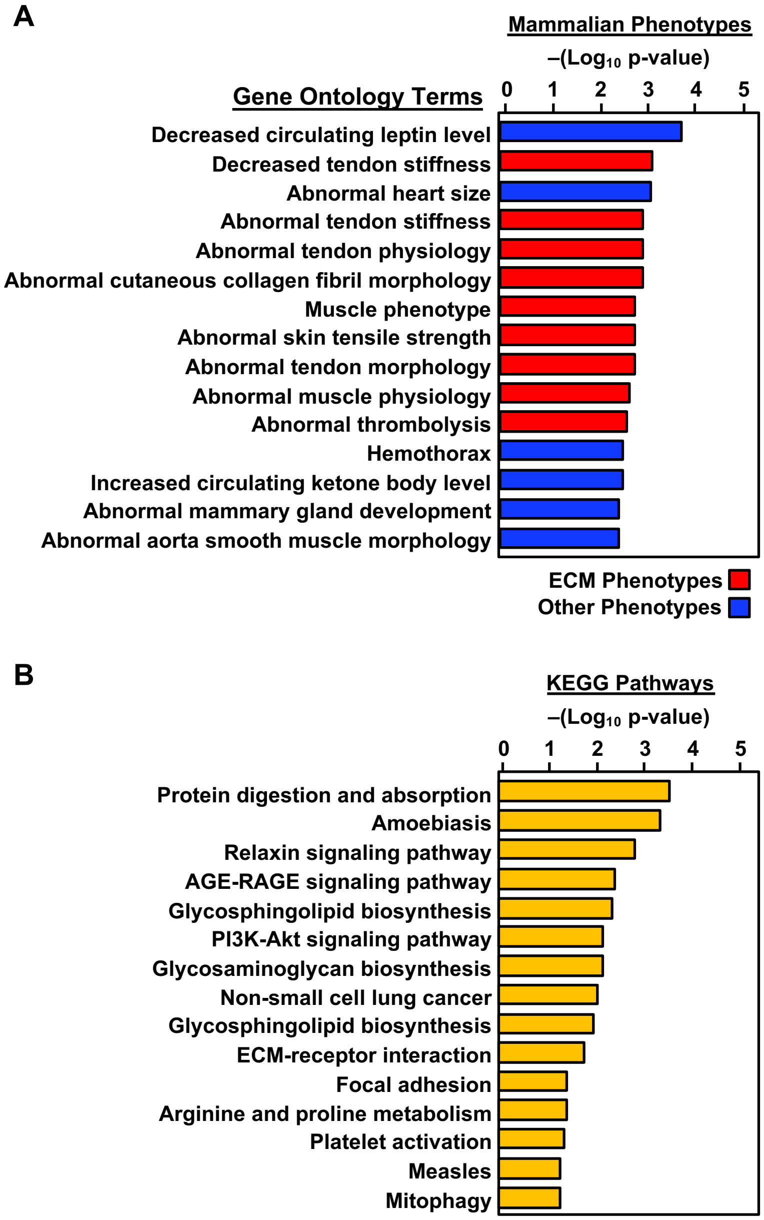


**Supplemental Figure S5. Prediction of mammalian phenotypes and pathways from genes associated with Rln-depleted ERα peaks**

Mammalian phenotypes and KEGG pathways for genes associated with Rln-depleted ERα peaks were evaluated using WEB-based Gene SeT AnaLysis Toolkit (WebGestalt) (58).

**(A)** Multiple gene ontology terms representing mammalian phenotypes related to extracellular matrix (ECM) were enriched.

**(B)** A set of KEGG pathways related to cellular signaling and metabolism were enriched.

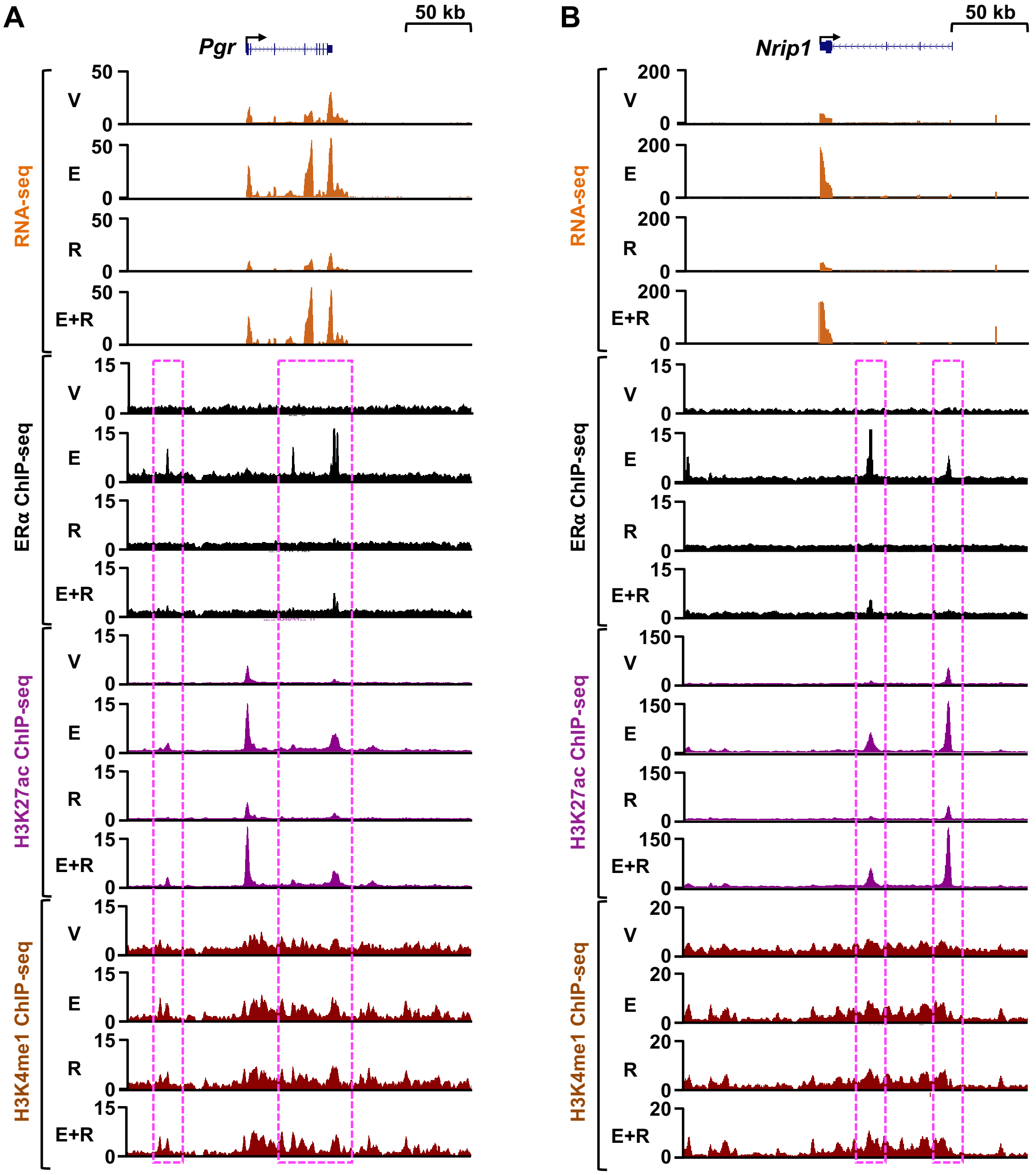


**Supplemental Figure S6. Genomic analysis of the effects of Rln treatment on genomic signaling by E2/ERα at classical estrogen-regulated genes.**
**(A and B)** Genome browser views of RNA-seq and ERα, H3K27, and H3K4me1 ChIP-seq data for the classical E2-regulated genes (A) *Pgr* and (B) *Nrip1* from ovariectomized mouse myometrium after treatment with vehicle (V), E2 (E), Rln (R), or both (E+R). Highlight (box with magenta dashed line) indicate regions of ERα enrichment from ChIP-seq. Gene schematic and scale bars are shown.

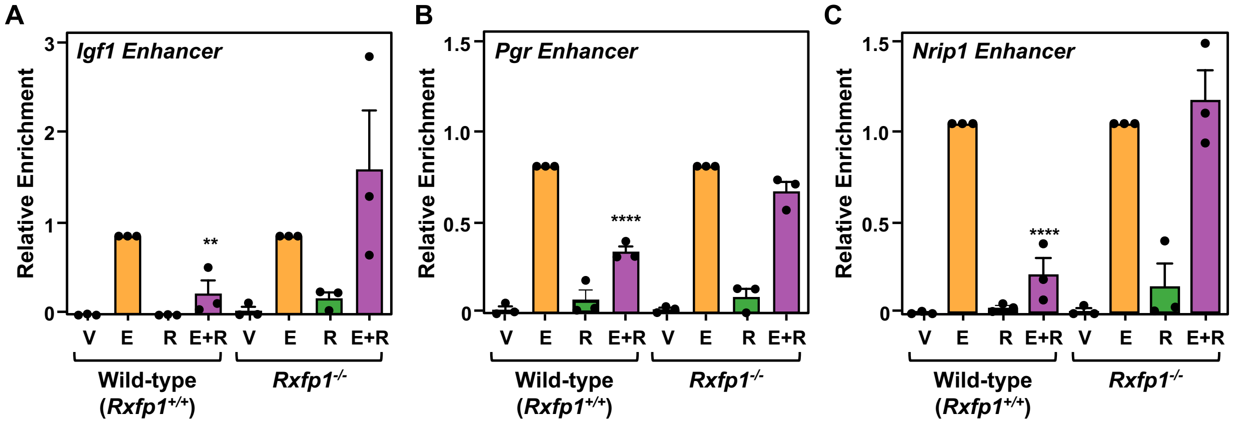


**Supplemental Figure S7. Genetic knockout of *Rxfp1*, the gene encoding the Rln receptor, abrogates the effects of Rln on the binding of ERα at multiple genome loci.**
**(A through C)** Binding of ERα in ovariectomized mouse myometrium, with or without *Rxfp1* knockout, at three well-characterized ERα enhancers upstream of (A) *Igf1*, (B) *Pgr*, and (C) *Nrip1* in response to various treatments, as determined by ChIP-qPCR. The results are shown as relative enrichment with respect to the E2-treated condition. Each bar represents the mean + SEM, n = 3, 10 mice/treatment. The asterisks indicate a significant difference from the corresponding control (V) (One-way ANOVA, $^{**}$ $p < 0.01$, $^{****}$ $p < 0.0001$).

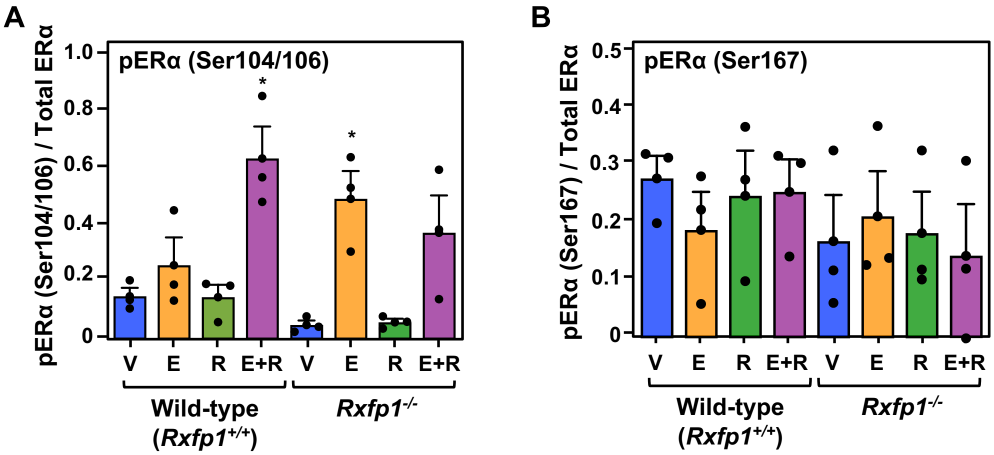


**Supplemental Figure S8. Effect of Rln on ERα phosphorylation in mouse myometrium with or without knockout of the gene encoding Rln receptor Rxfp1.**
**(A and B)** Densitometric quantification of ERα phosphorylation levels at Ser104/106 and Ser167 in myometrium from ovariectomized wild-type and Rln receptor knockout (*Rxfp1*$^{-/-}$) mice treated for 1 hour with the treatments indicated: vehicle (V), E2 (E), Rln (R), or both (E+R). Each bar represents the mean + SEM, n = 3, 5 mice/treatment. Asterisks indicate significant differences from the corresponding control (V) (One-way ANOVA, $^{*}$ $p < 0.05$).

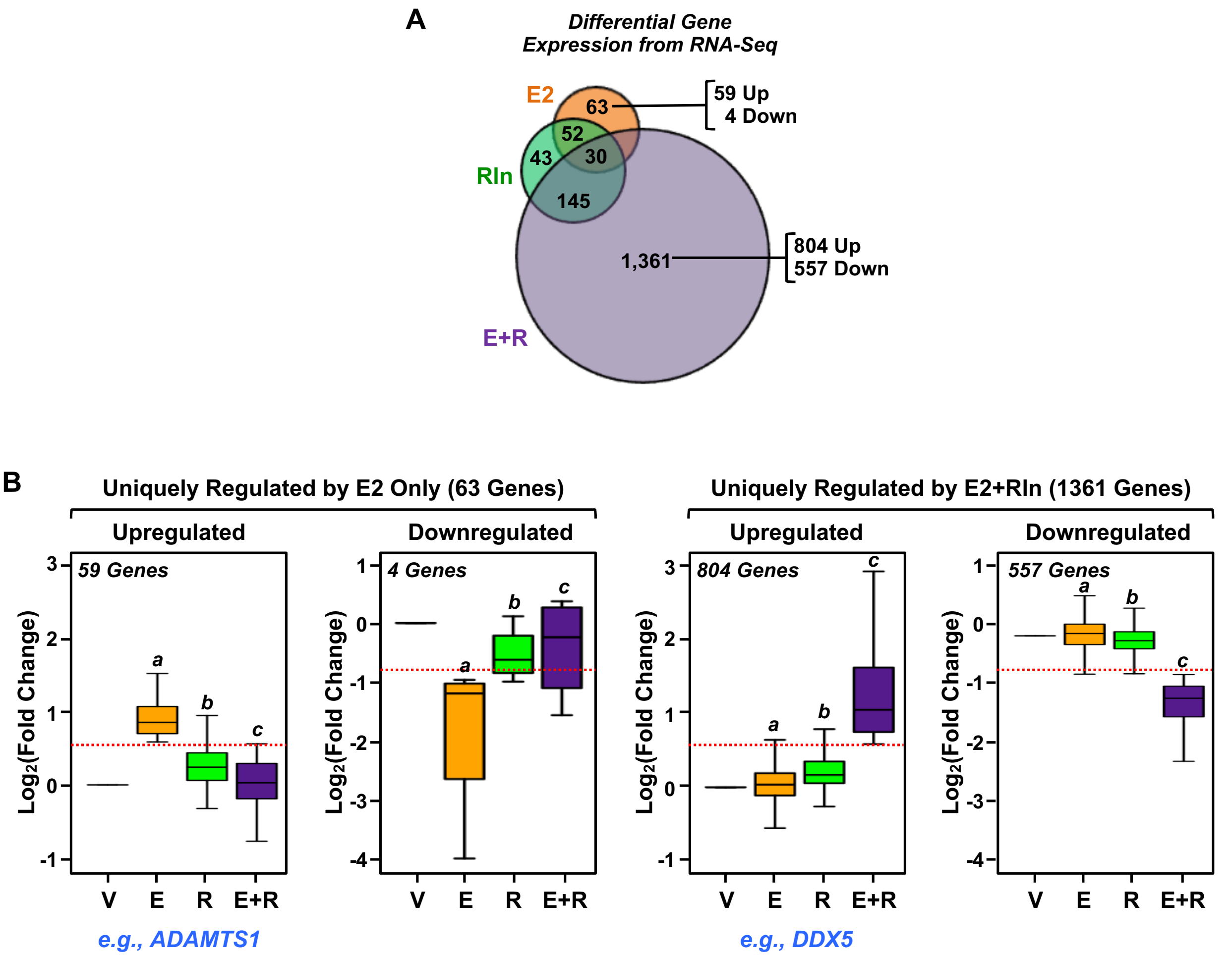


**Supplemental Figure S9. Genomic analyses of the transcriptional response to E2 and Rln in human myometrial cells.**
**(A)** Venn diagram showing differential gene expression from RNA-seq analyses in hTERT-HM-ERα cells after 1 hour of the indicated treatments. We categorized the treatment-regulated genes with a p-value filter [p value < 0.01 (q value < 0.05)] and fold change cutoff filter (≥ 1.5 or ≤ 0.67) versus vehicle. Only genes passing both filters were included in the category.
**(B)** Box plots showing the effects of the treatments on (*left*) genes uniquely regulated by E2 only (63 genes) and (*right*) genes uniquely regulated by E2+Rln (1361 genes) from the Venn diagram in (A). The data are expressed relative to control, which is set to 1. Dotted red lines indicate fold change cutoff ≥ 1.5 or ≤ 0.67 (log2 ≥ 0.58 or ≤ -0.79). Boxes marked with different letters are significantly different from each other and the control (Wilcoxon rank sum test, p-value < $2.2 \times 10^{-16}$). Genes listed below the "Upregulated" categories are representative of that category. (see browser tracks in Supplemental Figure S11, B and C).

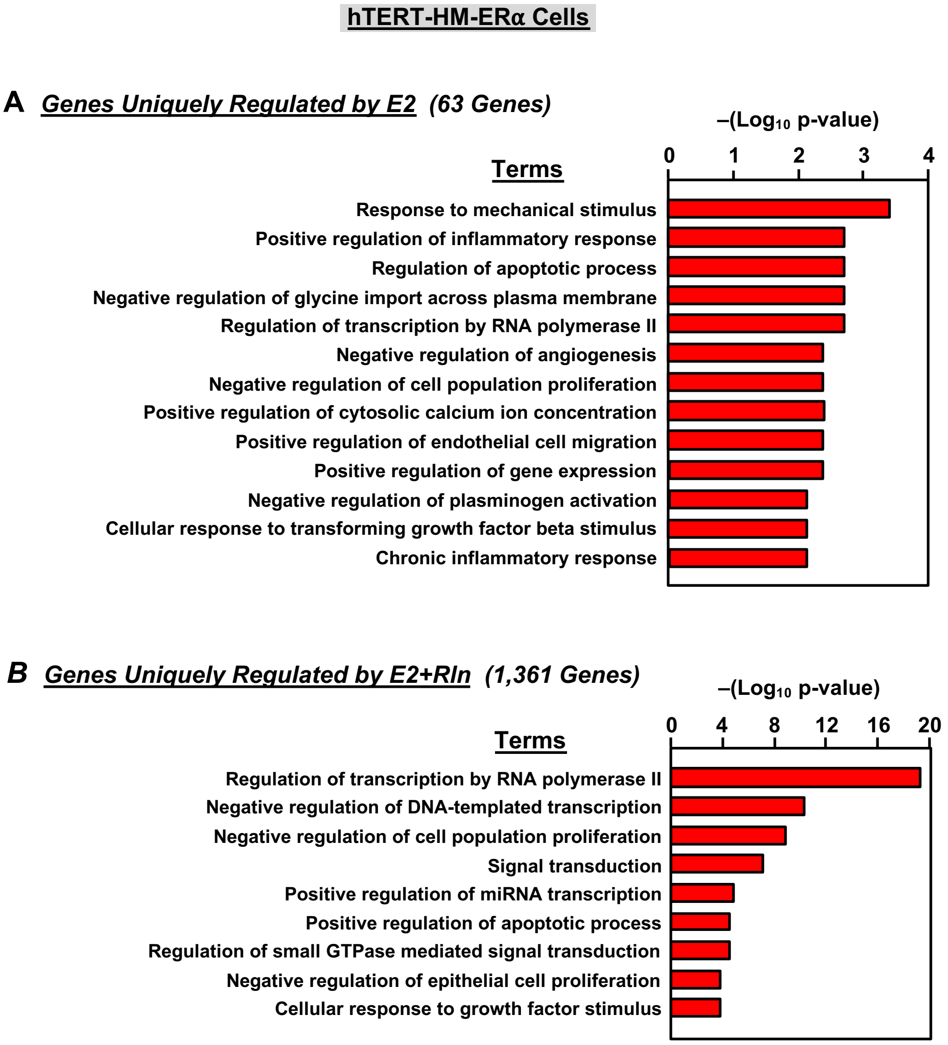


**Supplemental Figure S10. Gene ontologies for E2- and Rln-regulated genes in human myometrial cells.**
GO analyses of uniquely regulated genes from the Venn diagram in Supplemental Figure S9. The gene sets were evaluated using WEB-based Gene SeT AnaLysis Toolkit (WebGestalt) (59).
**(A)** GO analysis for genes uniquely regulated by E2 (63 genes).
**(B)** GO analysis for genes uniquely regulated by E2+Rln (1361 genes).

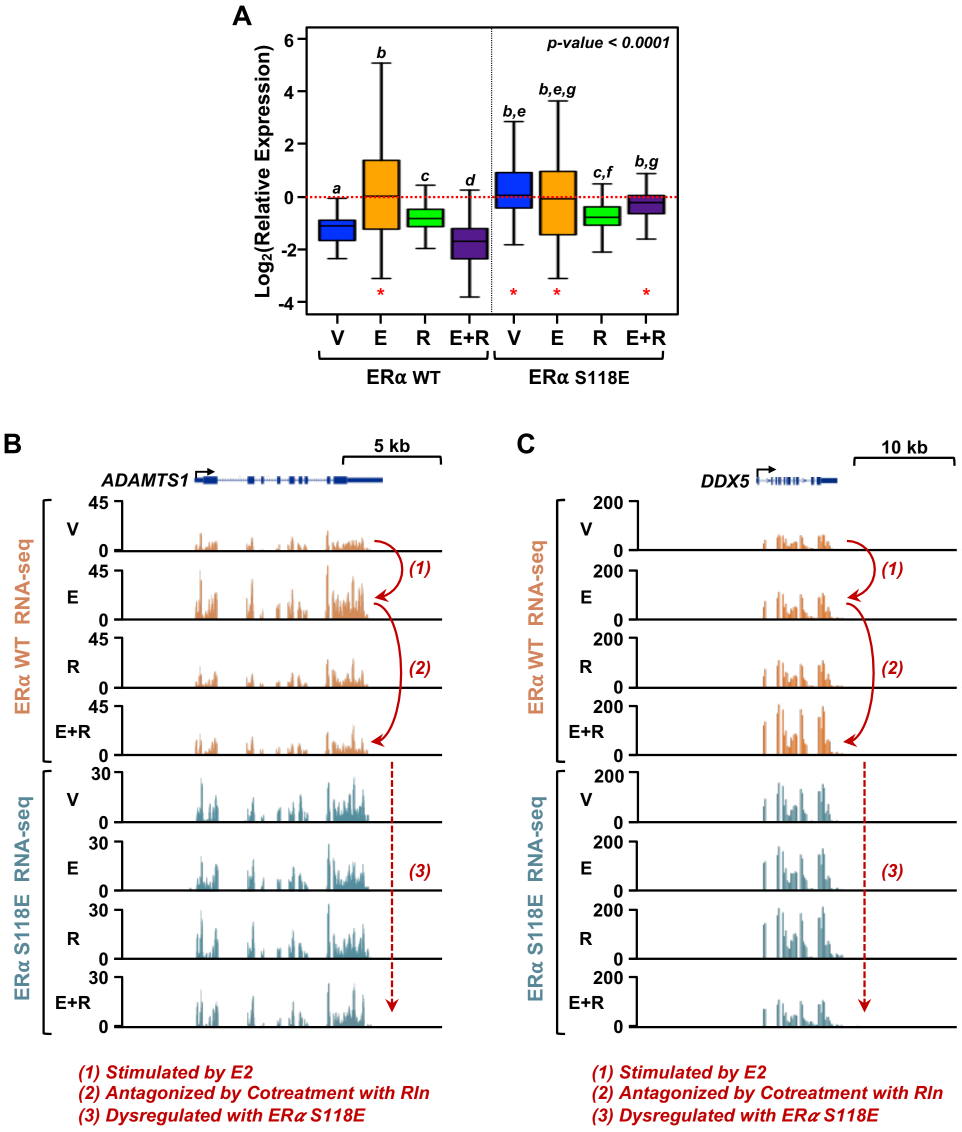


**Supplemental Figure S11. Genomic analysis of the effects of Rln treatment on genomic signaling by E2/ERα in human myometrial cells.**
**(A)** Box plots showing the effects of the treatments on genes upregulated by E2 and inhibited by cotreatment with Rln in hTERT-HM-ERα (WT or S118E) cells. The data are expressed relative to E2, which is set to 1. The E2 values, which were set to 1 for the analysis, were scaled to represent the range of values in the data set. Boxes marked with different letters are significantly different from each other (Wilcoxon rank sum test, p-value < 0.0001). The same gene set is represented for both WT and S118E ERα-expressing cells. Red asterisks indicate key comparisons discussed in the text.
**(B and C)** Genome browser views of RNA-seq data for representative genes from hTERT-HM-ERα (WT or S118E) cells after treatment with vehicle (V), E2 (E), Rln (R), or both (E+R). Gene schematic and scale bars are shown. The genes shown are (A) *ADAMTS1* (upregulated by E2 and inhibited by Rln) and (B) *DDX5* (further upregulated by E2+Rln). Labeled arrows indicate the types of regulation.

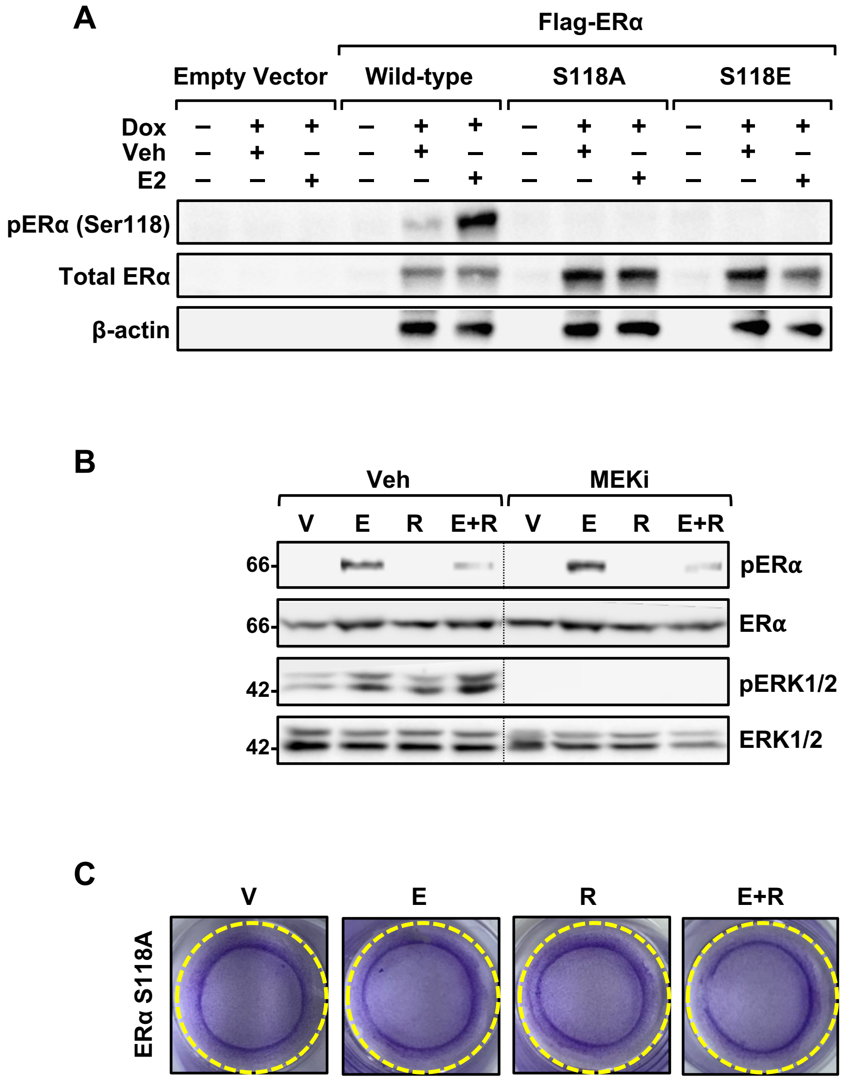


**Supplemental Figure S12. Mutation of ERα Ser118 to Ala inhibits E2-induced contraction of myometrial cells and inhibition of MAPK signaling does not prevent ERα Ser118 phosphorylation.**

**(A)** Western blotting of ERα phosphorylation levels at Ser118 in Dox-inducible hTERT-HM-ERα cells with wild-type, S118A, or S118E mutant ERα subjected to the treatments indicated for 30 minutes: vehicle (V), E2 (E), Rln (R), or both (E+R). β-actin serves as a loading control.

**(B)** Western blotting of ERα phosphorylation levels at Ser118 in Dox-inducible hTERT-HM-ERα cells treated with MEK inhibitor and vehicle (V), E2 (E), Rln (R), or both (E+R). β-actin serves as a loading control.

**(C)** Representative images of collagen gel contraction assays conducted in hTERT-HM-ERα cells with S118A mutant ERα subjected to the treatments as indicated in the Methods: vehicle (V), E2 (E), Rln (R), or both (E+R).

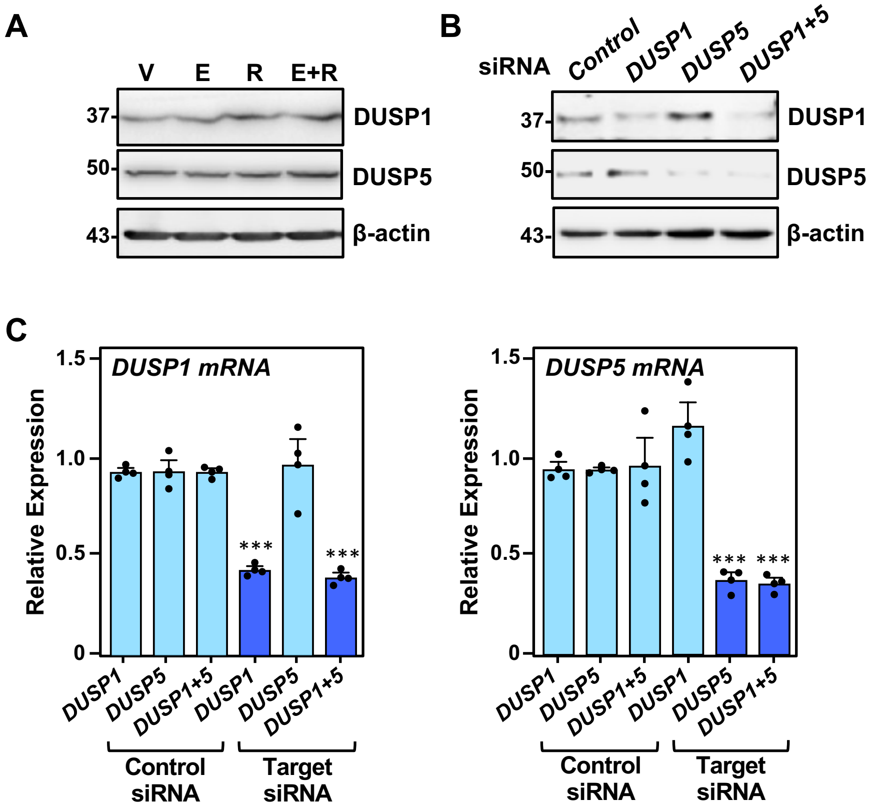


**Supplemental Figure S13. siRNA-mediated knockdown of *DUSP1* and *DUSP5* mRNAs.**
**(A)** Western blotting for DUSP1 and DUSP5 in hTERT-HM-ERα cells treated with vehicle (V), E2 (E), Rln (R), or both (E+R). β-actin serves as a loading control.
**(B)** Western blotting for DUSP1 and DUSP5 in hTERT-HM-ERα cells with siRNA-mediated knockdown of *DUSP1* mRNA, *DUSP5* mRNA or both. β-actin serves as a loading control.
**(C)** RT-qPCR analysis of the levels of *DUSP1* and *DUSP5* mRNAs in hTERT-HM- ERα cells subjected to siRNA-mediated knockdown of *DUSP1* mRNA, *DUSP5* mRNA or both, as indicated. Each bar represents the mean + SEM, n = 3. The asterisks indicate a significant difference from the corresponding control (One-way ANOVA, *** $p < 0.0005$).
